# DNA-segment-capture model for loop extrusion by structural maintenance of chromosome (SMC) protein complexes

**DOI:** 10.1101/325373

**Authors:** John F. Marko, Paolo De Los Rios, Alessandro Barducci, Stephan Gruber

**Affiliations:** Department of Molecular Biosciences and Department of Physics & Astronomy, Northwestern University, Evanston IL 60208 USA; Laboratory of Statistical Biophysics, Institute of Physics, School of Basic Sciences and Institute of Bioengineering, School of Life Sciences, Ecole Polytechnique Fédérale de Lausanne - EPFL, 1015 Lausanne, Switzerland; Center de Biochimie Structurale, INSERM, CNRS, Université de Montpellier, 34090 Montpellier, France; Department de microbiologie fondamentale, Université de Lausanne, 1015 Lausanne, Switzerland

## Abstract

Cells possess remarkable control of the folding and entanglement topology of long and flexible chromosomal DNA molecules. It is thought that structural maintenance of chromosome (SMC) protein complexes play a crucial role in this, by organizing long DNAs into series of loops. Experimental data suggest that SMC complexes are able to translocate on DNA, as well as pull out lengths of DNA via a “loop extrusion” process. We describe a Brownian loop-capture-ratchet model for translocation and loop extrusion based on known structural, catalytic, and DNA-binding properties of the *Bacillus subtilis* SMC complex. Our model provides an example of a new class of molecular motor where large conformational fluctuations of the motor ‘track’ - in this case DNA - are involved in the basic translocation process. Quantitative analysis of our model leads to a series of predictions for the motor properties of SMC complexes, most strikingly a strong dependence of SMC translocation velocity and step size on tension in the DNA track that it is moving along, with “stalling” occuring at subpiconewton tensions. We discuss how the same mechanism might be used by structurally related SMC complexes (*E. coli* MukBEF and eukaryote condensin, cohesin and SMC5/6) to organize genomic DNA.

## I. INTRODUCTION

A fundamental problem faced by all living cells is dealing with the fantastic length of their chromosomal DNAs - millimeters in bacteria, and centimeters to meters in many eukaryote cells. Given the ≈ 300 base pair (bp) statistical segment length of double-stranded DNA (dsDNA), the ≈ 5 × 10^6^ bp chromosomes of bacteria and the ≈ 10^8^ bp chromosomes of mammalian cells are in the realm of extremely long and flexible polymers. Left to their own devices, the multiple chromosomal DNAs of these lengths found in bacteria and in eukaryote cell nuclei should become highly entangled with one another [1–3]. Nevertheless, cells are able to completely separate their chromosomal DNAs from one another, and to keep them from being entangled with one another.

Two enzymatic machines are believed to play an essential role in the topological simplification of chromosomal DNAs *in vivo*. The first are type II topoisomerases (topos), which are enzymes which pass one dsDNA through a catalytically generated break in a second dsDNA [4]. Following strand passage, type II topos reseal the dsDNA break and then release the DNA strands, changing knotting or linking topology [5] (Fig. 1a). The key enzyme of this type in bacteria is called topo IV, while in eukaryote cells it is topo II*α*. Type II topos are ATPases, and use stored energy to complete their reaction cycle. Although type II topos are known to have the capacity to channel energy released by ATP hydrolysis into simplification of entanglement topology of DNAs to levels below that expected in thermal equilibrium [6–8], this effect is insufficient by itself to eliminate entanglement of whole chromosomes in cells.

**FIG. 1:**
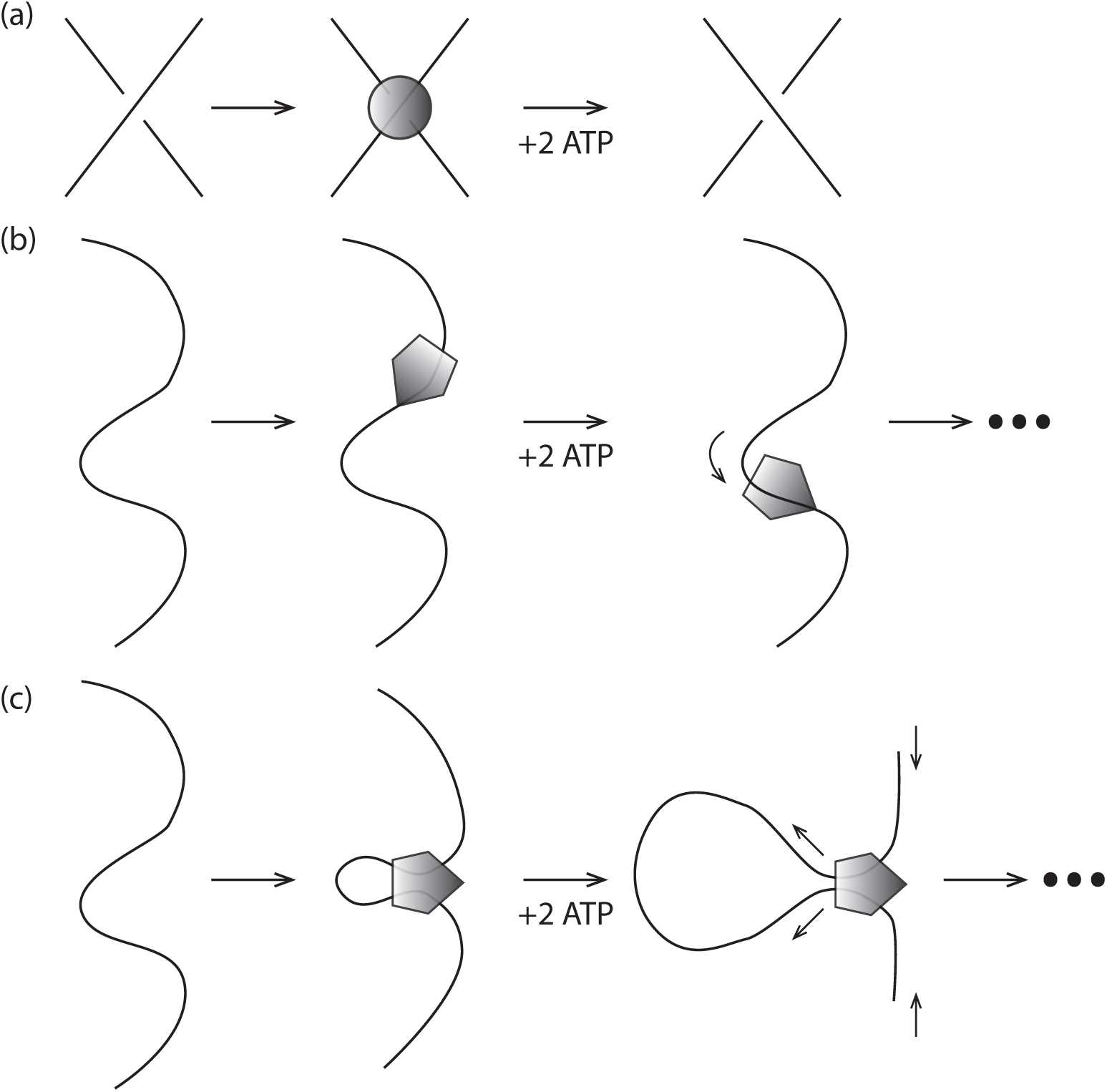
Two active (ATP-hydrolysis-powered) protein machines responsible for DNA topology control *in vivo*. (a) Type II topoisomerases are able to catalyze the transfer of a dsDNA segment through a second piece of dsDNA strand by generating a transient dsDNA break (not shown). Two ATPs are bound and hydrolysed per double-strand DNA exchange. (b) SMC complexes are able to translocate along DNA. (c) SMC complexes are thought to be able to generate the gradual growth of size of DNA loops through active ‘loop extrusion’. Note that only the first step of translocation and loop extrusion is shown; repetition of ATP binding and hydrolysis of additional ATPs can lead to processive enlargement of DNA loops.

A second class of enzymatic machine which appears essential to topological simplification of whole chromosomes are Structural Maintenance of Chromosome (SMC) complexes, which are also ATP-consuming molecular machines (Fig. 1b) [9–12]. SMC complexes are based on ≈50-nanometer (nm) long coiled-coil protein domains that close into ring-shaped structures (Fig. 2). An array of experimental data point to the capacity of SMC complexes to translocate along DNA in a directed fashion, as well as to mediate DNA “loop extrusion” processes, whereby an initially small DNA loop is increased in size processively. SMC complexes are thought to be involved in a number of DNA-organizing processes, including tethering of the two halves of bacterial chromosomes together [13–17], DNA-sequence-nonspecific compaction of chromosomes via crowding together of DNA loops [3, 18, 19], and also the bringing together of specific signal sequences spaced by distances of 10^5^ to 10^6^ bp along DNA molecules in specific orientations [20, 21]. Recent single-molecule experiments on yeast condensin have directly observed translocation [22] and loop-extrusion [23] functions.

**FIG. 2:**
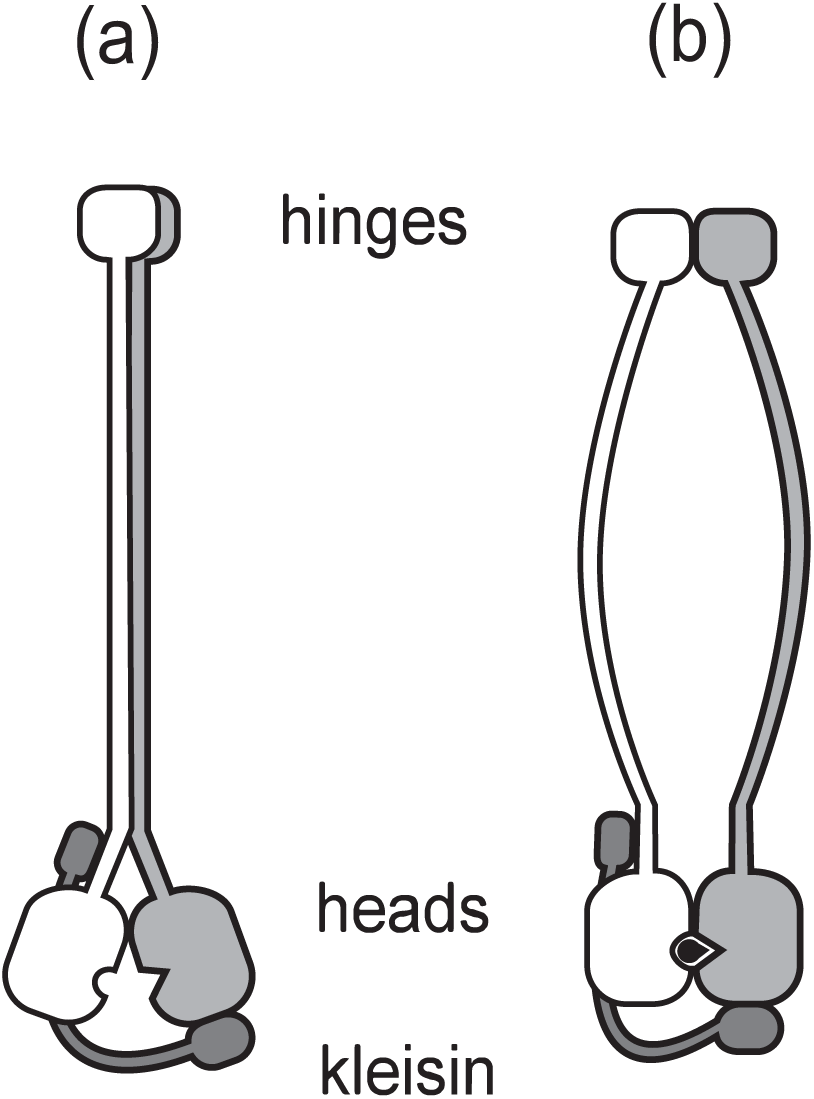
Schematic view of SMC complex, showing the long (≈ 50 nm) coiled-coil domains which are linked at a ‘hinge’ domain and which have ATP-binding ‘head’ domains, and additional “kleisin” protein subunits which are able to bridge the head domains (dark gray). In the ATP-unbound (apo) state (a) the long coiled-coils associate (left). ATP (black) binding allows the head domains to bind together and change the conformation of the coiled-coils (b). ATP hydrolysis and release of hydrolysis products returns the enzyme to its apo state (a).

At their core, these functions involve organization of long dsDNAs into series of loop structures, plausibly facilitating detection of DNA connectivity and control of DNA topology. Impressively, SMC complexes appear to be an ancient class of enzymes, with structurally similar orthologs occuring in all kingdoms of life. Key SMC complexes under current study include the bsSMC complex in the bacterium *Bacillus subtilus*, the MukBEF complex found in the bacterium *E. coli*, and the condensin, cohesin and SMC5/6 complexes found in eukaryote cells.

A major puzzle regarding SMC complexes is their DNA translocation and loop-extrusion mechanisms, given that they are structurally unrelated to known DNA-based protein machines, e.g., RNA polymerases, DNA polymerases and DNA helicases. Structural and enzymatic analyses of the bsSMC complex have led to a model whereby small (≈ 600 bp) loops of DNA are captured and incorporated together to gradually extrude loops [24]. Here, we analyze this model from a quantitative theoretical biophysics point of view. Although we focus on bsSMC, much of our modeling appears to be applicable to eukaryote condensin, which has been the focus of a great deal of recent attention.

In Materials and Methods we begin by describing the sequence of conformational states thought to occur for bsSMC during its ATP binding, hydrolysis, and product release cycle, and then we describe how DNA conformational changes can be coupled to these protein conformational states so as to accomplish translocation of a SMC along a dsDNA molecule. An essential ingredient of the model is the capture of thermally generated DNA loops, which leads to a strong dependence of the translocation rate on tension in the dsDNA ‘track’, illustrating its novelty as a molecular motor system where the flexibiity and deformation of the motor track plays an essential role.

As shown in Results our model is sufficiently biophysically based that we can make predictions about the transition rates between the different SMC/DNA states and therefore the cycling and translocation dynamics of the SMC/DNA combination. In Results we also describe a few plausible schemes whereby DNA loops could be extruded by SMC complexes: in one, two SMC translocators are coupled together, but in the others we show how a single SMC complex may be able to generate processive DNA loop extrusion, as observed in experiments on yeast condensin [23]. In the Discussion section we review the key results of the paper, and predictions of the model for specific experiments.

## II. MATERIALS AND METHODS

### A. Model for SMC translocation along DNA

While we will focus on the specific case of bsSMC (where there is at present the best structural understanding), the model we now construct will be general enough to be applicable to all SMC complexes. Fig. 2 shows the ‘body plan’ of bsSMC: like other SMCs they consist of long coil-colied proteins (called SMC proteins) which dimerize at one end at “hinge” domains, and which have ‘head’ domains at their other ends of Walker type which bind and hydrolyse ATP. The two heads are able to bind together when they bind ATP, and ATP hydrolysis releases this binding.

All SMC complexes also contain an additional protein, often called a “kleisin” subunit, which bridges the two heads using two distinct interfaces, breaking the symmetry of the SMC dimer (the two SMC proteins are a homodimer in the bacterial case, and a highly symmetric heterodimer in the eukaryote case). Various species of SMC have additional subunits which likely play key roles in DNA targeting and regulation of the complex, but for this paper we will focus on the interplay between the hinge-coiled-coil-head SMC proteins and the bridging protein.

### B. Structural states of SMC complexes

SMC complexes can be found in three distinct nucleotide-binding states, either the “apo” state with no ATP bound, the ATP-bound state, or the ADP-bound state [24]. *In vivo*, ATP concentration is essentially fixed, which indicates that these three states define a chemical reaction cycle: the apo state (Fig. 3a) will bind free ATPs (Fig. 3b), then hydrolyse them (Fig. 3c), and then release the hydrolysis products to return to the apo state (Fig. 3a). While there may be additional states that a particular SMC complex might be able to occupy, all SMC complexes are thought to go through this ATP binding/hydrolysis cycle in vivo.

**FIG. 3:**
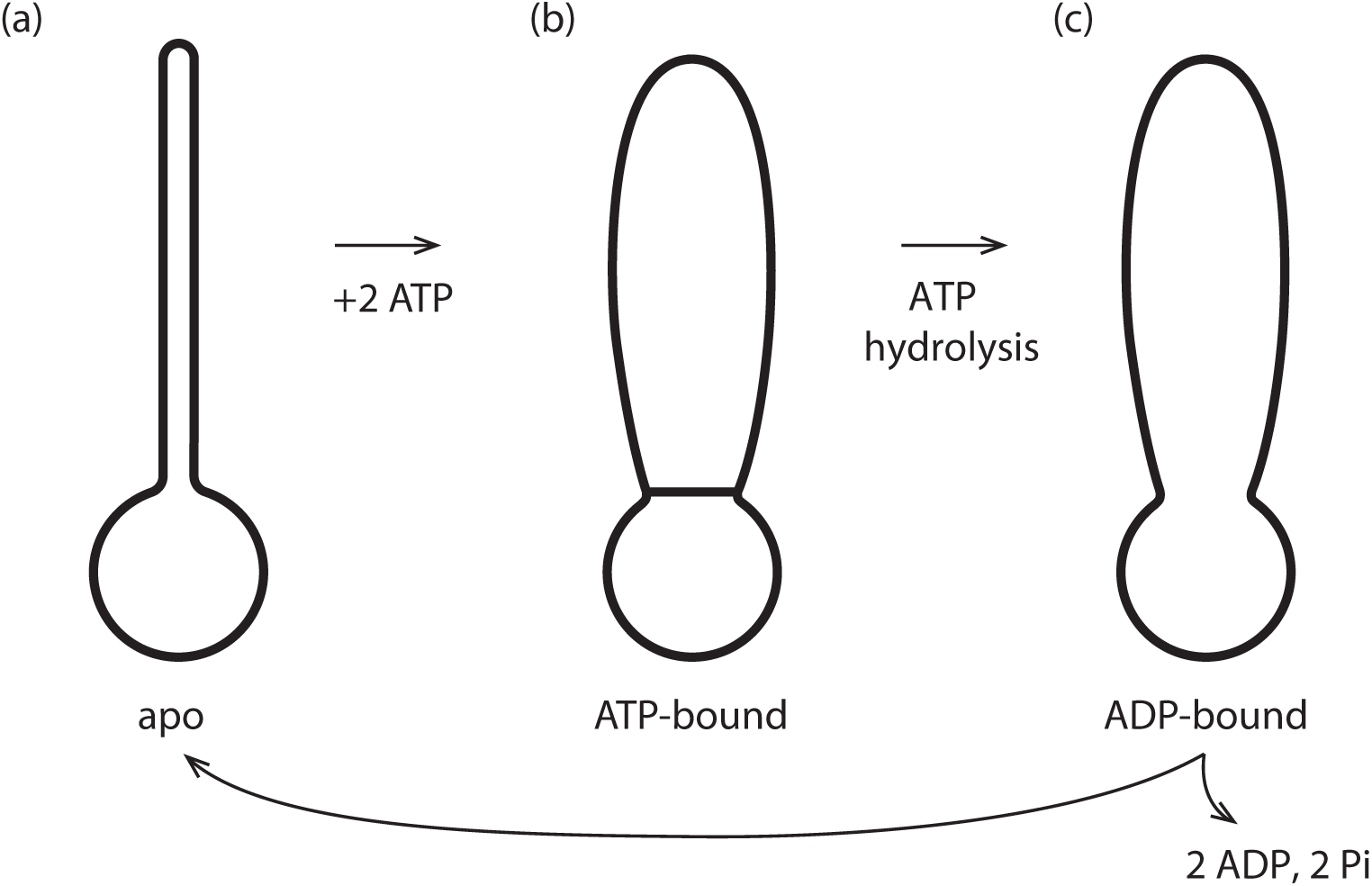
SMC protein complex states. (a) The apo (non-nucleotide-bound) state has a rod-shaped conformation with its two coiled-coil domains bound together and a small open ‘lower’ compartment; (b) ATP binding opens the two coiled-coil domains resulting in the formation of a larger ‘upper’ compartment; (c) ATP hydrolysis opens the ‘gate’ between the upper and lower compartment. Release of ADP leads to a reset of the enzyme to the apo state of (a).

For the case of bsSMC, it is established that these three ATP-binding states correspond to different structural states of the enzyme. In the apo state, the long coiled-coil arms of the bsSMC complex are stuck together. At the end of the coiled coils, the heads are also held together, and with the bridging protein define a single ring-like ‘lower compartment’ at the base of the complex (Fig. 3a). Then, when ATP binds, there can be a conformational change in the complex which bows the two coiled-coil arms apart, opening a second “upper compartment”. The protein “wall” between the two compartments is held closed by the ATPs bound between the two “head” subunits (Fig. 3b); in effect the ATPs act as ‘molecular glue” which holds the two heads together while facilitating the opening of the upper compartment.

When the ATPs are hydrolysed, the wall between the two compartments is opened, and they merge into one large compartment (Fig. 3c). Then, when the ADPs are released (the product of ATP hydrolysis), the two coiled-coils bind together, closing the top portion of the compartment, and returning the complex to the apo state (Fig. 3a).

In Fig. 3, the (a) to (b) transition involves two distinct steps: nucleotide binding, followed by protein conformational change. We also note that under conditions where ATP and ADP are present at appreciable concentration, there will be competition for binding. Similarly, the transition from (b) to (c) also can be resolved into two successive steps of ATP hydrolysis followed by conformational change. In the next section we will take account of this finer picture of the dynamics of the complex; we will also consider all reverse steps, allowing a complete thermodynamic accounting of energy use by the reaction cycle.

### C. Reaction cycle for translocation of SMC along DNA

Our model is sketched in Fig. 4a in terms of a reaction cycle coupling the ATP-driven conformational changes of the SMC complex with DNA binding and unbinding. We show the same states using molecular models for bsSMC in Fig. 4b (based on Ref. [24]). The upper diagrams are intended to make molecular connectivity and topology clear; gray circles indicate noncovalent DNA-protein interactions. The lower diagrams illustrate practical realizations of molecular conformational changes based on available experimental information [24].

**FIG. 4:**
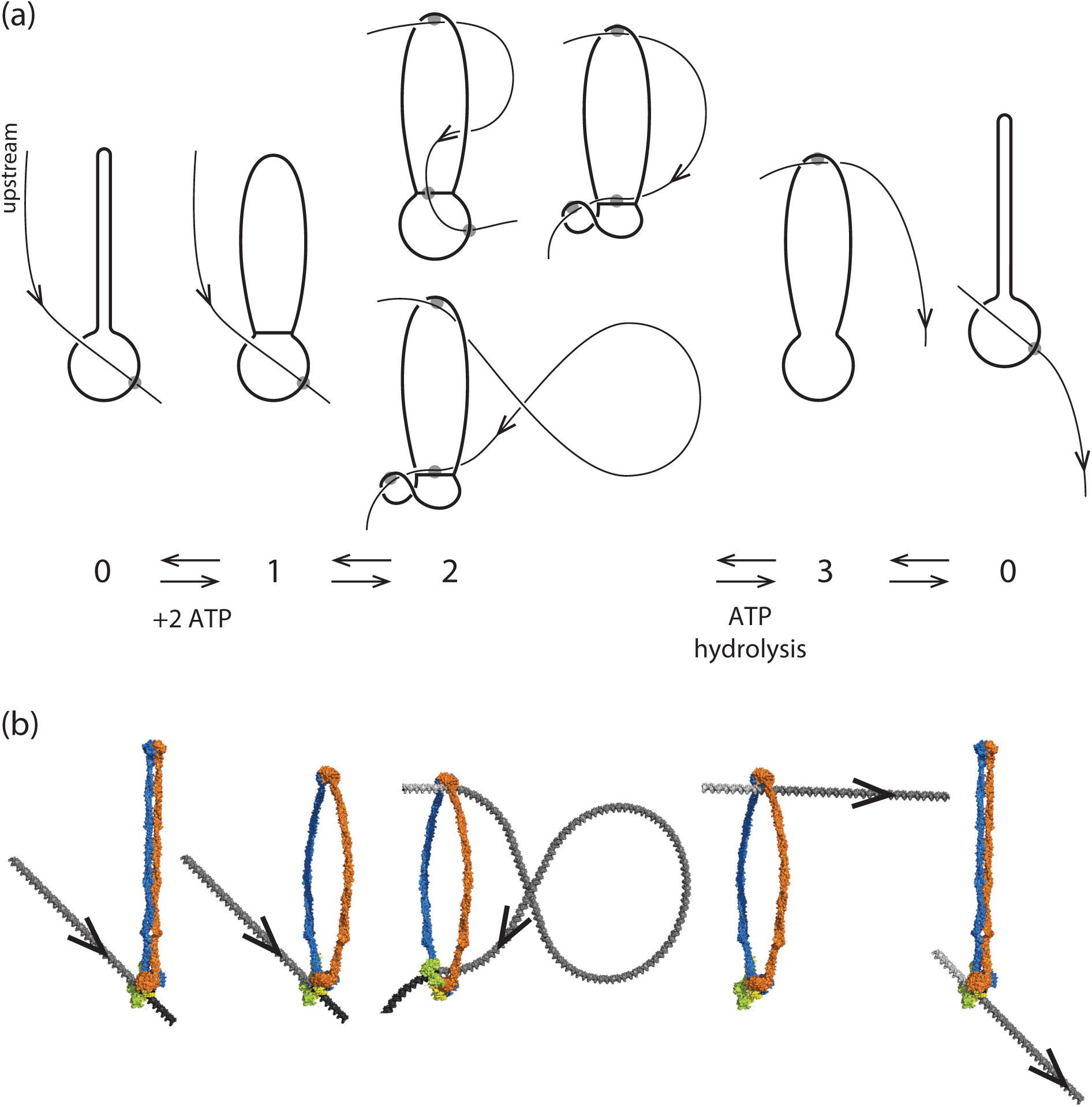
Reaction diagram for SMC reaction with DNA that can generate translocation. (a) Schematic/topological view of reaction, which starts with the SMC complex in the apo state 0 and with DNA bound in the lower compartment. Binding of ATP permits SMC head engagement and opening of the upper compartment (1), allowing a DNA loop to be captured in it (2). Then, ATP hydrolysis opens the wall between the two compartments, allowing the loop to pop out, transferring DNA contour length to the right (3). The coiled-coils then bind back together, pushing the DNA back into the lower compartment, returning the SMC-DNA complex to state (0), but translocated on the DNA by approximately the contour length captured in the loop in state (2). Three sketches of state 2 are shown, corresponding to a purely topological picture (state 2, top left), where the SMC complex lower chamber is bent to facilitate DNA loop capture (state 2, top right), and where a large DNA loop with a crossing is captured (state 2, bottom) (b) Molecular structure diagram (based on data of Ref. [24]), showing corresponding states of SMC complex and DNA. Note the reorientation of the DNA segment by the “folding” of the lower compartment of the SMC complex in state 2, similar to the lower sketch of Fig. 4a. DNA structures with appropriate curvature were generated with the online tool 3D-DART and assembled together with manually fitted models of Smc-ScpAB in PyMol.

The reaction cycle begins and ends with the enzyme in its apo state “0” (Fig. 4a, leftmost and rightmost states). A DNA segment is encircled by the lower compartment in bsSMC and yeast cohesin in the apo state [25, 26], either sterically held, or possibly bound non-covalently to a DNA binding site as observed in eukaryotic SMC complexes [27–30]. The configuration of state 0 can be imagined to be the result of the type of ‘loading’ reaction associated with initiation of SMC complex activity.

An essential feature of this initial state is the direction of “threading” of the DNA through the lower compartment, plus DNA-protein binding, which breaks symmetry “upstream-downstream” or “left-right” along the DNA (the “upstream” DNA “to the left” of the binding site is labeled “upstream” in state 0 of Fig. 4a and carries an arrow throughout the reaction cycle of Fig. 4a). This breaking of symmetry is expressed via DNA binding to a site (Fig. 4a, state 0, gray circle) displaced from the SMC complex centerline, with DNA axis constrained to point in an oblique direction (the general case given the broken symmetry of the protein-DNA complex). Importantly, this broken symmetry dictates that rotation of the state 0 complex by 180*^◦^* around the coiled-coil “hairpin” does not return the complex to a similar conformation. In turn, this breaks symmetry between DNA positions upstream and downstream from the SMC lower compartment. For bsSMC we suppose that there is a mechanism for firmly holding DNA in the lower compartment: this might be an enthalpic interaction, or due to the small size of the lower compartment the interaction could be steric for the case of bsSMC.

In apo state 0, there is continuously binding and unbinding of ATP to the two SMC binding sites, and in the conformational state 0, when two ATPs are bound, there is a possibility of a transition to the SMC conformation where the two Walker ATPases are stably bound together (Fig. 4a, state 1). We suppose that concomittant opening of the two coiled-coils exposes DNA binding sites in the inside of the upper protein “compartment”, permitting capture of a second DNA site as a result of thermal fluctuation of the enzyme complex and the double helix (Fig. 4, transition to state 2). Capture of a second DNA site *in cis* will be more likely than *in trans* because of the large local concentration of sites along the same DNA molecule.

From structural studies, it is known that there is a highly conserved DNA binding site at the bottom of the upper compartment, near the Walker ATPase “heads” [25, 31, 32]. In addition, there is a DNA-binding site at the top of the upper compartment (near the hinge domain) conserved across different SMC family members [24, 33, 34], which seems dispensable in the case of bsSMC [25]. We presume that in the apo state, the closed coiled-coils prevent binding of DNA to these hinge and head binding sites. A number of SMC complexes have been observed to be able to bind to DNA so as to trap small DNA loops, in the range from 70 nm to 200 nm in size (200 to 600 bp) [35–38]; this likely involves having DNA bound at two of these sites spaced appropriately to facilitate DNA distortion. These pieces of experimental data are all in accord with our model for the 0 *↔* 1 *↔* 2 pathway.

Capture of the DNA in the SMC protein upper compartment is key to the mechano-chemical cycle of our model. For directional translocation to occur, the DNA segment captured by the upper compartment must be preferentially to one side or the other of the initially bound DNA site. In Fig. 4a, this preference is to the “upstream” direction travelling opposite to the arrowhead relative to the initially bound DNA site. There must be such a preference, due to the broken symmetry of the DNA-SMC complex: this is a result of its chirality, of the direction that the DNA is initially “threaded” through and bound the lower compartment, and of the asymmetry in the conformational fluctuations of the complex.

Three possible configurations for state 2 of Fig. 4a illustrate how SMC complex conformational changes may bias loop capture to one side of the initially bound DNA site. “Folding” of the lower compartment (state 2, upper right configuration) can align the DNA molecule so that it is “pointed” towards the SMC upper compartment, making loop-capture of upstream DNA more likely than downstream DNA.

A variant of this (Fig. 4a, state 2, lower configuration) shows a larger DNA loop of “teardrop” shape with a DNA crossing captured by the complex, a conformation again facilitated by the “folding” of the lower SMC compartment. The interaction shown at the top of the SMC protein compartment in Fig. 4 could be anywhere in the compartment, and might even be a weak, transient interaction of the DNA loop “trapped” inside the upper compartment.

After capture of the second upstream DNA site, ATP hydrolysis can occur, releasing the Walker ATPases from one another, opening the “gate” between the two compartments, and eliminating the DNA binding sites in the lower compartment. The DNA loop is then free to relax, leading to state 3 of Fig. 4, where the DNA is now held only in the upper compartment, at the upstream position established by the preceding DNA loop capture step.

Finally, ADP release and conformational change of the SMC back to the apo state pushes the DNA out of the upper compartment, leading to it to being rebound by the lower compartment. Due to the asymmetry of state 0 and consequent asymmetry of DNA loop capture, the protein complex has translocated upstream on the DNA, or equivalently the DNA has been “pumped” through the SMC protein, by a distance comparable to the contour length of the captured DNA loop.

As in all molecular motor models, the cycle is driven in one direction (rather than diffusing back and forth) by the directionality of ATP binding and hydrolysis. The tendency for translocation to occur in one direction along the DNA is determined by the asymmetry between binding upstream rather than downstream DNA in the transition from state 1 to state 2. This asymmetry follows from the broken symmetry between upstream and downstream DNA in state 0 of Fig. 4a.

### D. Kinetic-thermodynamic model

To have directional translocation along the DNA, the SMC complex must capture a second site *in cis* preferentially to one side of the initially bound site. This can occur for the DNA-SMC complex in state 1 for thermodynamic reasons: the conformation of the state-1 in general must have a free energy difference between capture of a second DNA site to one side or the other, because of the broken symmetry of the SMC complex (the two states where DNA “to the left” versus DNA “to the right” is captured are distinct and not related by a symmetry operation of the protein-DNA complex, e.g., by a rotation).

Given this asymmetry, there must be a difference in rate of trapping DNA to the left or to the right: the model of Fig. 4 presumes that the segment to the left is always trapped, but of course one could imagine that there could be a “backwards” step caused by capture of a segment “to the right”. As long as there is a bias to capture a segment to one side or the other (the general case), repeated cycles in one direction (determined by the directionality of ATP hydrolysis) on average will generate translocation in a specific direction along the DNA, defined by the direction in which the DNA is threaded through the lower compartment in state 0.

One must also account for binding, hydrolysis and release of two nucleotides per reaction cycle, along with consideration of ADP binding in the apo state, and the possibility of reversed reactions (ATP synthesis), while maintaining consistency with thermodynamics (the kinetics must be consistent with the free energy released by ATP hydrolysis). Without the directional drive of the ATP hydrolysis step, even with the asymmetric loop binding, our reaction would cycle in both directions at equal rates, leading to diffusion rather than translocation. We now describe a more complete model (Fig. 5a) and its mathematical formulation, and we show how it can be reduced to a four-state model of the form of Fig. 4.

**FIG. 5:**
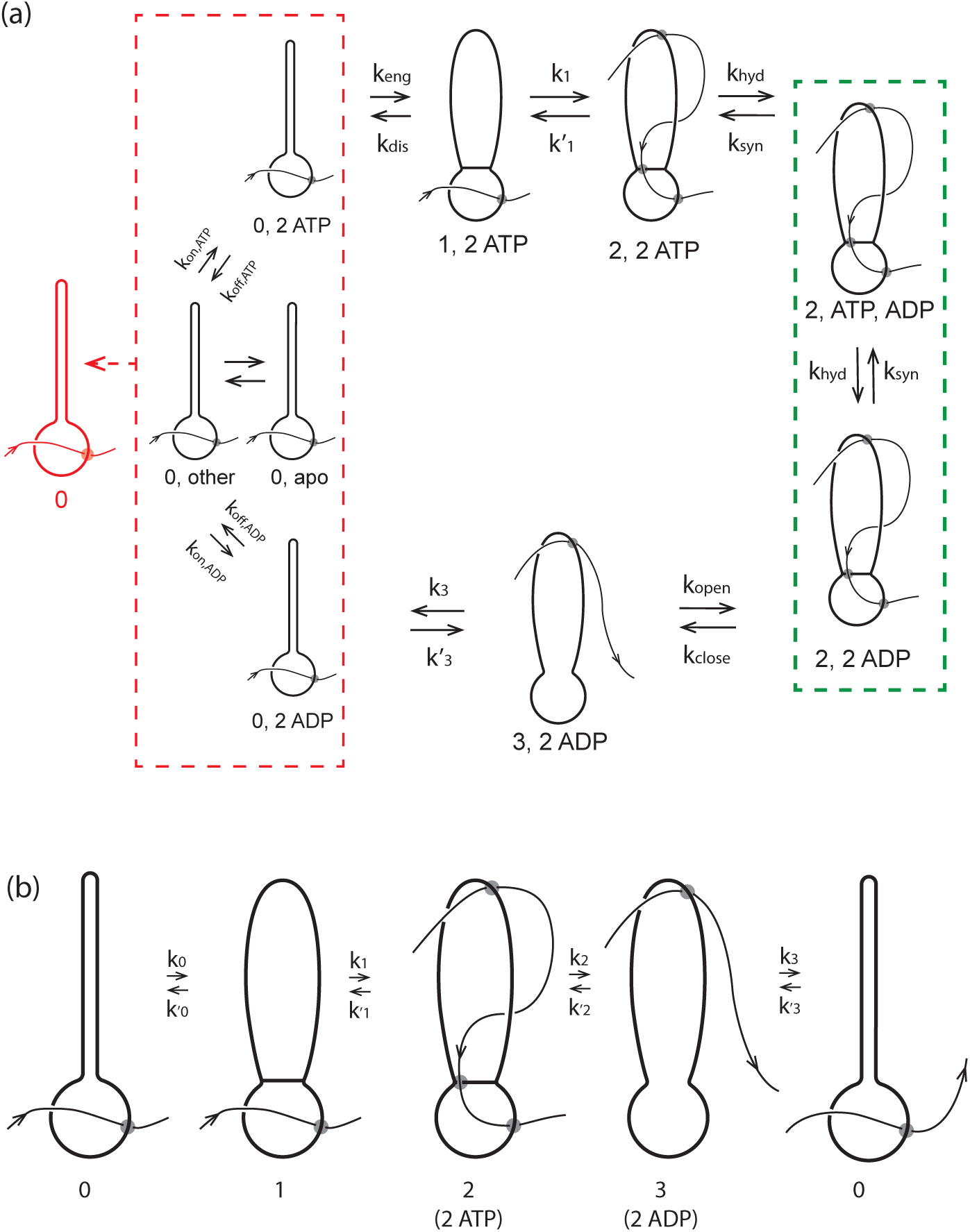
Complete reaction diagram for SMC translocating along DNA. (a) Complete reaction diagram starts with the SMC complex in state 0 and with DNA bound in the lower compartment, and able to bind nucleotides (ATP or ADP: left red dashed box includes 16 nucleotide-binding states; “other” represents the 13 states other than apo, 2 ATPs bound, and 2 ADPs bound). State labels are of the form N,n where N is the state number and n is bound nucleotide state (e.g., 0,2ATP refers to conformational state 0 with 2 ATPs bound). Binding of 2 ATPs permits SMC head engagement and opening of the upper compartment (state 1), allowing a DNA loop to be captured in it (state 2). ATP hydrolysis opens the wall between the two compartments, allowing the loop to pop out, transferring DNA contour length to the right (state 3). Finally the coiled-coils bind back together, pushing the DNA back into the lower compartment, returning the SMC-DNA complex to state 0, translocated on the DNA by approximately the contour length captured in the loop in state 2. (b) Coarse-grained model, where the different nucleotide-binding states are combined together to form state 0 (left boxed states in (a)), and where ATP hydrolysis and loop release processes are combined together into a single 2-3 transition (the 2,ADP state of (a) is effectively “’integrated out”; state 2 now represents 2,ATP).

In Fig. 5a, each state is labeled by the conformation state (0, 1, 2 or 3 as in Fig. 4) followed by nucleotide state (e.g., 2ATP means two ATPs bound; ATP,ADP means one ATP and one ADP bound, and so on). The dashed lines group related states together for reasons which will become apparent shortly. Fig. 5 has less topological detail than Fig. 4 (crossings of DNA over/under protein are not emphasized), but the topology of the SMC-DNA states (0, 1, 2 and 3) of Fig. 5 correspond to those of Fig. 4.

#### 1. 0↔1: ATP binding and SMC head engagement

We begin with the apo state, which is able to bind and release ATP, and ADP if it is present. We consider all 16 possible nucleotide combinations (no nucleotide bound, ATP bound, ADP bound, or ADP plus inorganic phosphate bound at each of the two binding sites). Because nucleotide binding/unbinding will occur rapidly in the apo state (the binding sites are exposed) our strategy will be to lump all those states together, into the “0” conformational state (Fig. 5a, left red dashed box). The nucleotide occupation state will be treated as being pre-equilibrated, which makes sense given the rapidity of nucleotide exchange relative to the slower conformational and hydrolysis reactions.

We presume that only in the state where two ATPs are bound (Fig. 5a, 0,2ATP state) can the SMC complex undergo the transition from the rod-like state 0 to the open-coil state 1, where the two ATP-charged heads are engaged (“stuck”) together. The net rate for head engagement will depend on the probability of two ATPs being bound, which will in turn depend on the concentration of ATP (and ADP if sufficiently large). Of course there is also a “disengagement” reverse rate, accounting for the possibility that thermal fluctuations break the two ATP-charged heads apart.

The mathematical details of the description of this transition are along the lines of standard binding kinetics models [39] and along with known ATP affinity data for bsSMC and other ATPases [40, 41] are discussed in Supplementary Data.

#### 2. 1↔2: Reversible DNA loop capture

Once the upper compartment is open, two DNA binding sites are exposed, allowing a DNA loop to be captured (Fig. 5a, state 2). As discussed above, due the left-right asymmetry of the protein complex (all SMC complexes possess this asymmetry), we can expect a loop to form “from the left” via capture of upstream DNA or “from the right” with unequal rates. Since the DNA “track” is unpolarized, so that left-right symmetry-breaking of the SMC complex is essential to ensure processive translocation. We suppose that capture of an upstream DNA loop dominates, and we ignore “looping from the right” (this could be included but will not essentially change the model). DNA loop formation of the transition from state 1 to state 2 is reversible, since state 2 is held together by non-covalent bonds and can be destabilized by tension in the DNA.

The forward rate *k*_1_ requires a loop of DNA to be formed, and thus depends on DNA mechanics (its persistence length *A* = 50 nm, or the slightly shorter effective persistence length of nucleoid-associated-protein-coated DNA [42, 43] as found in a bacterial cell). Any tension in the DNA, *f*_DNA_, also will decrease *k*_1_ by penalizing large loops [44] and as a consequence slowing down the loop-capture rate. Two key points emerge from this transition: the step size for translocation will be on the order of the size of the DNA loop that is trapped by the SMC (the position of the second, upstream interaction site between DNA and SMC); and, increased DNA tension will reduce the step size. Again, there is the possibility of a reverse transition whereby the DNA loop is released: this reverse transition is likely to be increased in rate by increasing DNA tension.

Our mathematical description of the loop-capture kinetics is based on established results from the DNA-looping and cyclization literature [44, 45]. Details and parameters can be found in Supplementary Data.

#### 3. 2↔3: ATP hydrolysis, phosphate release, and loop release

When the ATPs are hydrolysed, the two heads are released from one another, and the wall between the upper and lower compartments is opened. This requires two successive hydrolysis steps with two intermediates with one and two ADPs bound (2,ATP,ADP and 2,2ADP of Fig. 5a, respectively, inside the green dashed box). We take the forward rate *k*_hyd_ = 20 sec*^−^*^1^ [40]. Despite being exceedingly slow under almost any conceivable experimental condition, we will also consider reverse ATP synthesis steps.

ATP hydrolysis causes the two Walker ATPase heads to dissociate from one another, releasing DNA from the bottom of the upper compartment. We assume that this step also eliminates the assumed DNA binding capacity of the lower compartment. The DNA site bound in step 0 is released, allowing the DNA to pop through the now-opened “gate” between the lower and upper compartments (Fig. 5a, state 3). This is the translocation step, since it moves DNA contour (arrow) that was on the left side of the SMC complex in state 0, to the right side of the SMC.

The result is the 2-ADP-bound state 3, which has DNA held only in the upper compartment (Fig. 5a, state 3,2ADP); this state is likely to be highly transient since the ADPs have relatively low affinity and are not being held in place by the engaged heads. In steady state we can eliminate the two ATP hydrolysis steps to obtain a single pair of rates linking conformational state 2 to state 3.

The DNA loop capture and release process involves formation of DNA crossings, which can generate supercoiling of the DNA template. However, our model does not require crossings of a specific sign, and by forming crossings of opposite signs, supercoiling of the DNA can be avoided. On the other hand, for SMC complexes that trap DNA loops of specific or preferred chirality, it is conceivable that supercoiling of the DNA template could be generated along with translocation. The details and parameters used in our computations are described in Supplementary Data.

#### 4. 3↔0: ADP release and conformational reset

The final step of the reaction cycle is the closing of the upper compartment (reformation of the rod-like conformation of the SMC coiled-coils) which resets the SMC back to conformational state 0 and pushes the DNA back into the lower compartment where it rebinds. Relative to the initial state, the SMC complex is now translocated along DNA by an amount of approximately *ℓ*, the size of the loop captured during the 1→2 transition. The rate of this process is given by *k*_3_ = 10 sec^−1^. The rate 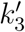 for the reverse of this process is actually now determined by the free energy released by ATP hydrolysis during the cycle, as discussed in Supplementary Data.

### E. Reduction to the four-state model

We can simplify the model of Fig. 5a to a four-state cycle corresponding to the four major conformational states described earlier (Fig. 4). We take advantage of the rapid equilibration of nucleotide-binding states in SMC state 0 (Fig. 5a, left red dashed box) and consider them together, with net probability *P*_0_. Given pre-equilibration of nucleotide binding, the transition from the block of 0 states to state 1 occurs only when two ATPs are bound. Note that *k*_0_ connects ATP concentration to forward cycling: *k*_0_ increases linearly as [ATP] is increased from zero, and saturates at high [ATP].

A further simplification is combining together ATP hydrolysis and conformational transitions by eliminating the ATP hydrolysis/synthesis step, thus combining the 2,ATP,ADP and 2,ADP,ADP states into a single, composite state. (Fig. 5a, green dashed box). This is possible if we restrict ourselves to consideration of steady-state cycling of our model. Formulae for the resulting steady-state rates *k*_2_ and 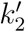 connecting the 2,2ATP state to the 3,2ADP state are derived and described in detail in Supplementary Data.

Once the model is expressed as shown in Fig. 5b, in terms of the four SMC-DNA conformational states of Fig. 4. The four-state model is exactly equivalent to the detailed model of Fig. 5a (as long as we consider only steady-state cycling); it allows us to focus on the experimentally relevant DNA-tension-dependence associated with loop capture in *k*_1_ and 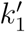 and the ATP concentration dependence in *k*_0_.

### F. Reversibility, energy consumption, and thermodynamic consistency

Our model has the feature that all forward and reverse chemical rates are defined, meaning that it is microscopically reversible, and has a thermodynamic interpretation. In turn this indicates that the free energy liberated by ATP hydrolysis must be equal to the free energy dissipated during the reaction cycle. This constraint is used to set the value of the rate 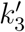 (see Supplementary Data, Energy consumption during the reaction cycle).

Thermodynamic consistency leads to the desirable feature that our reaction cycle runs forwards only when there is free energy to be gained from ATP hydroysis (i.e., for sufficiently large [ATP], dependent on ATP/ADP chemical equilibrium [46]). Our model makes nontrivial predictions for the dependence of the cycling rate on concentrations of each of ATP, ADP and phosphate (Supplementary Data, Free energy release from ATP hydrolysis and equilibrium nucleotide concentrations).

### G. Steady-state cycling rate

We can compute the steady-state probabilities *P_i_* of each of the states of Fig. 5b, and the steady-state cycling and DNA translocation rates. In steady state there must be equal flux into and out of each state, giving the set of equations

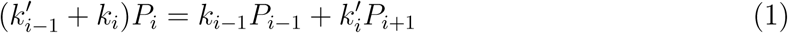

where the subscripts are considered modulo 4, and where state 2 represents the 2,2ATP state (as in Fig. 5b). In addition we have the constraint that the total probability of all states sums to unity, or *P*_0_ + *P*_1_ + *P*_2_,_2ATP_ + *P*_2_,_ATP_,_ADP_ + *P*_2_,_2ADP_ + *P*_3_ = 1.

This system of equations is analytically solvable, although rather lengthy and uninformative formulae are obtained. It can be shown that the flux (cycling rate *k*_cycle_) through the cycle (any of 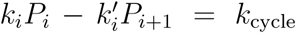 for *i* = 0*, · · ·*, 3) is proportional to ([ATP][ADP]_eq_[Pi]_eq_)^2^*/*([ATP]_eq_[ADP][Pi])^2^ *−* 1, i.e., that nucleotides and phosphates must be perturbed from their equilibrium values for a cycle (nonequilibrium) to occur (see Supplementary Data). The cycling rate is straightforward to compute from the exact steady-state equations (1).

### H. Translocation rate

The cycling rate for the SMC-DNA complex *k*_cycle_ is not yet the translocation rate. The forward velocity of the SMC along its DNA template is given by *ℓk*_cycle_, but if there is a load-a force *f*_load_ impeding the SMC’s progress along DNA of *f*_load_ (Fig. 6a) - one can expect some slippage of the SMC backwards along the DNA. Notably, for simple translocation, the load can be considered as a parameter at least partially independent of the tension in the DNA template *f*_DNA_, allowing description of slippage as a process separate from the loop-capture-translocation. The simplest model is one where there is slippage of the DNA from one of its binding sites at some point during the translocation cycle.

**FIG. 6:**
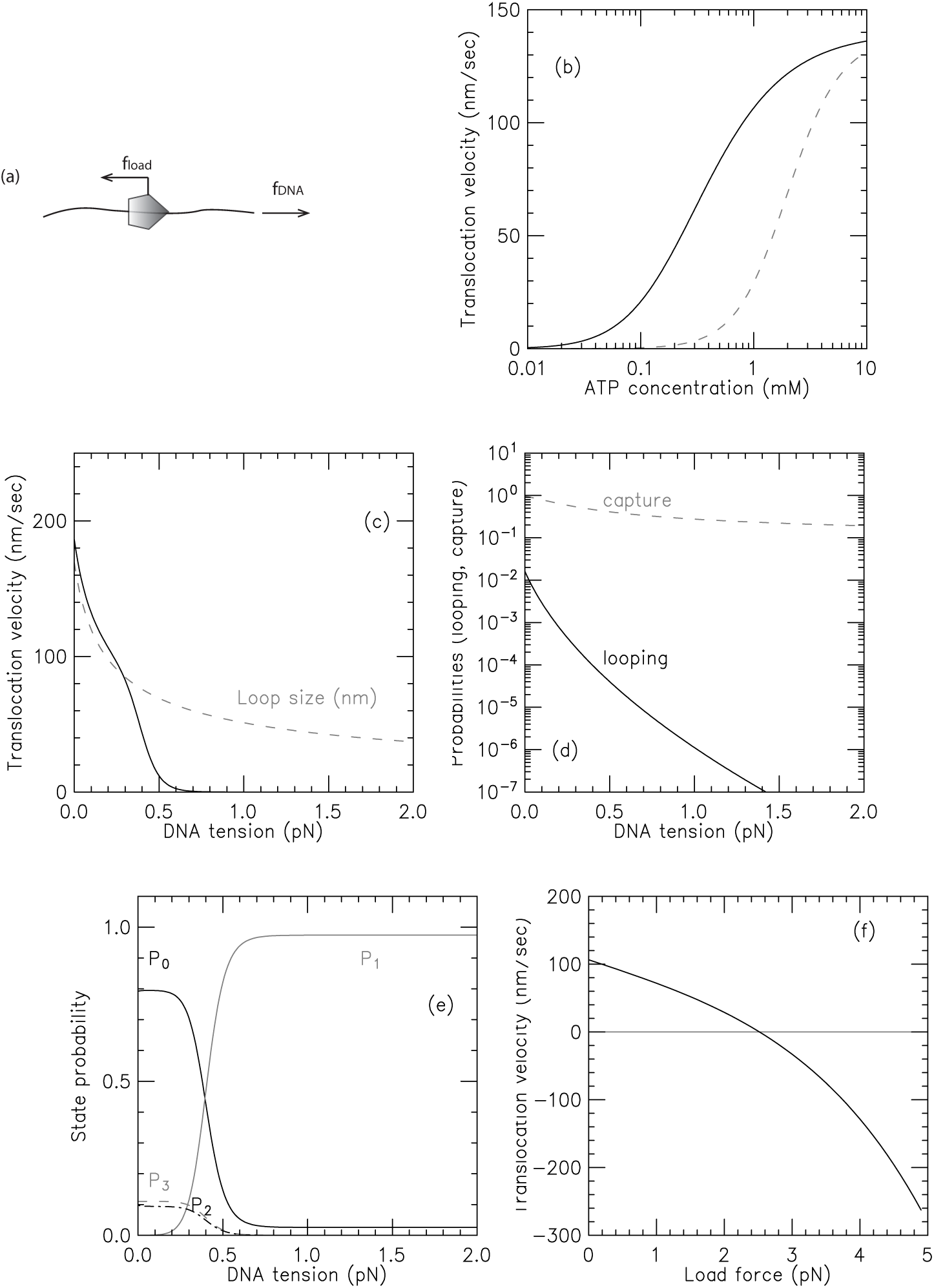
Translocation for SMC DNA loop-capture model dependence on experimental control parameters. (a) Distinction between tension in the DNA versus load force applied directly to the translocating SMC complex. (b) Solid curve: velocity as a function of ATP concentration for zero load and DNA tension of *f*_DNA_ = 0.2 pN. Dashed curve: result of same model in the presence of ADP; the velocity-ATP curve is simply shifted to larger ATP concentration values on the log [ATP] scale. (c) Velocity as a function of DNA tension for zero load and ATP concentration of 1 mM shown by solid black curve; gray dashed curve shows the loop size. For the parameters chosen, DNA tension strongly (exponentially) suppresses SMC translocation for forces in excess of approximately 1 pN, but never reverses the direction of translocation. (d) Variation of the loop-formation Boltzmann factor and the loop-capture probability with DNA tension, corresponding to (c) (see Supplementary Data for formulae). (e) Probability of the four states of the model of Fig. 4, as a function of DNA tension, corresponding to (c). (f) Velocity as a function of load force for template tension of 0.2 pN and ATP concentration of 1 mM. For the parameters chosen, the load force reverses the direction of the SMC complex along DNA for forces in excess of ≈ 3.5 pN.

We suppose that such events are associated with a slipping length Δ and that the barrier opposing these slip events is on the order of *ε*_slip_. A simple force-driven barrier-crossing model gives a reverse slipping velocity of

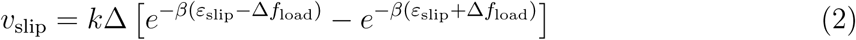

where the two terms describe random slip in the load force direction and against it; at *f*_load_ = 0, slipping in either direction is equally (im)probable, giving *v*_slip_ = 0. The constant Δ is the range of deformation of the DNA-SMC bound state that is required to break their chemical contact, likely of nm dimensions; here we take Δ = 2 nm. We take the slipping energy *ε*_slip_ = 14*k_B_T*, which is the work that must be done to break the DNA-SMC contact by force. Given this slip velocity, the net translocation velocity is

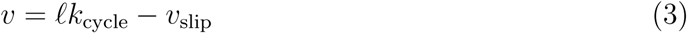

## III. RESULTS

### A. Dependence of translocation velocity on DNA tension and load force

Since the translocation velocity is observable in single-molecule SMC-DNA experiments, we examine its behavior for reasonable choices of parameters. The solid line in Fig. 6b shows the (exact) translocation velocity as a function of ATP concentration at zero load force and DNA tension of *f*_DNA_ = 0.2 pN, showing how the translocation velocity saturates at high ATP concentration. For the solid line, ADP concentration is held fixed at 1 *µ*M; the dashed line shows how translocation is slowed when ADP is increased to 0.1 mM, due to the decreased free energy drive associated with the increased ADP concentration (see Supplementary Data).

Fig. 6c shows the translocation velocity as a function of template DNA tension *f*_DNA_ and for fixed ATP concentration of 1 mM, fixed ADP concentration of 1 *µ*M, and zero load force, showing how translocation is slowed down by tension in the DNA being translocated along, and then is finally stopped. This results from the decrease of the rate of loop formation due to increased tension (see Supplementary Data). The translocation crucially depends on the rate *k*_1_, which is a product of the loop-formation Boltzmann factor and the loop capture probability (see Supplementary Data). Fig. 6d shows how these two quantities vary with DNA tension for the same ATP and ADP concentrations; the probability for loop formation dramatically decreases (Fig. 6d, solid curve), shutting off translocation. There is also a weaker suppression effect (Fig. 6d, gray dashed curve) generated by the decrease in loop size (Fig. 6c, gray dashed curve).

Fig. 6e shows the (exact) evolution of the probabilities of the four states as DNA tension is increased (for the same ATP and ADP concentrations as in Fig. 6c-d): at low tension, state 0 is highly probable due to the conformational change rate being rate limiting. Then, as force is increased, DNA loop capture becomes rate-limiting, resulting in a decrease of *P*_0_ and an increase of *P*_1_. For the parameters used here, the probabilities of *P*_2_ and *P*_3_ are similar; they start at 5 to 10% at low force and then are both pushed to zero when the tension in the DNA becomes large enough to stall the motor, enforcing occupation of state 1 with high probability.

Finally, Fig. 6f shows the translocation velocity as a function of load force, which eventually not only slows the SMC translocation velocity but can reverse it, pushing the enzyme back by inducing a high rate of slipping events. The relatively large stall via load force may be difficult to observe, given that most experimental arrangements to apply a load force will also apply DNA tension, which will quench translocation at a low force. It should be kept in mind that it will be difficult in practice to entirely decouple load force and DNA tension, since pulling on the enzyme will also to some degree pull on the DNA.

Key predictions of our model follow from the loop-capture mechanism and are apparent from Fig. 6c. First, stalling of translocation should occur for subpiconewton forces, well below the few-pN stall forces typically observed for cytoskeletal ATP-powered motors such as kinesins. Second, the average step size of SMCs should be reduced by increased DNA tension (the average translocation step size should approximately correspond to the loop size shown in Fig. 6c). Third, the DNA tension cutoff for translocation with DNA tension should be adjustable by changing SMC complex size (more precisely, by adjusting the size of DNA loops that can be captured). It may be possible to adjust the cutoff *l*_0_ (or perhaps the DNA loop-shape dependent constant *D*) by changing the length of the SMC coiled-coils, with longer coils leading to a cutoff of translocation at lower forces, and with shorter coils leading to a cutoff at higher forces.

In Supplementary Data we include additional data for the extrusion velocity ignoring slippage, showing dependence on ATP/ADP ratio (Supplementary Data Fig. 1), total nucleotide concentration (Supplementary Fig. 2), versus force for varied total nucleotide concentration (Supplementary Data Fig. 3), and for varied phosphate dissociation constants (Supplementary Data Fig. 4). We also include data for the probabilities of the four states as a function of ATP/ADP ratio (Supplementary Fig. 5) and DNA tension (Supplementary Data Fig. 6).

### B. Approximation of small reverse rates and rapid head opening: Michaelis-Menten-like behavior

In an experimentally-relevant limit, the complete model of Fig. 5 (numerical steady-state results of Fig. 6) (1), has a relatively simple analytical solution which is physically informative. Given the parameters described above and for ATP concentrations in the physiological range (near mM), the reverse rates of Fig. 5b are negligible relative to the forward ones, with one exception, namely 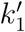 as we now explain.

The transition from state 1,ATP back to state 0,ATP is strongly suppressed by the free energy favoring head engagement; ATP synthesis is much slower than ATP hydrolysis when ATP and ADP are anywhere near mM concentrations, the transition from the opened state 3,2ADP back to 2,2ADP is supressed by the free energy cost of forcing the gate closed in the absence of ATP, and the transition from 0,ADP to 3,ADP is similarly supressed by the cost of forcing the “hairpin” open. The only transition which can have backward steps that can compete with forward steps under physiologically-relevant conditions is the DNA-loop-capture transition from 1,ATP to 2,2ATP (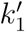 of Fig. 5). The balance between these forward and reverse transitions can be changed using DNA tension, making this step crucial to controlling the cycling rate. In fact, eliminating all of the reverse steps except 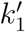 leads to nearly identical results to the model with all reverse steps, and also allows a simple analysis of the model’s behavior.

We can also make the approximation that little time is spent in the 2,ADP,ADP and 2,2ADP states relative to the 2,2ATP state, which results from the head opening rate (*k*_open_) being fast compared to the ATP hydrolysis rate (*k*_hyd_). Given the relatively slow rate of ATP hydrolysis, this is likely, and allows the further approximation that *P*_2_ *≈ P*_2_,_ATP_; in our approximate treatment we take *P*_2_ = *P*_2_,_ATP_.

Given these approximations (*P*_2_ = *P*_2_,_ATP_, all reverse rates set to zero except 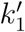, see Supplementary Data) the cycling rate is

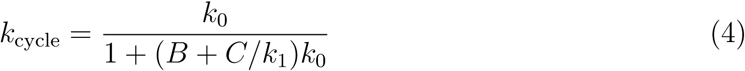

where *B* = 1*/k*_2_ + 1*/k*_3_ and 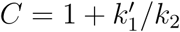 are used to write the simplified final expression. We note that *k*_0_ and *k*_1_ are easily related to experimental control parameters; *k*_0_ is controlled by ATP concentration, and *k*_1_ is controlled by the tension in the DNA (*f*_DNA_).

The cycling rate has a Michaelis-Menten form with respect to *k*_0_: for low *k*_0_ (achieved for sufficiently low ATP concentration), the cycling rate approaches *k*_0_, but for sufficiently large *k*_0_, the cycling rate saturates at *k*_cycle_,_max_ *→* 1*/*(*B* + *C/k*_1_), reflecting that transitions other than 0*→*1 are rate-limiting for the reaction cycle when *k*_0_ is sufficiently large (*i.e.*, *k*_0_ *» k*_0_*_,m_* = 1*/*[*B* + *C/k*_1_]).

For fixed *k*_0_ (ATP concentration) the behavior of *k*_cycle_ with *f*_DNA_ (via *k*_1_) is interesting. As the tension is increased above *f*_0_ ≈ 0.1 pN, *k*_1_ starts to drop, and eventually *k*_1_ will become rate-limiting, with 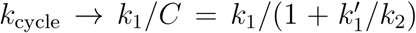. We note that this effect, of tension in the template (*f*_DNA_) controlling the rate of motion of a motor, is rather unusual. We also note that *f*_DNA_ is not, strictly speaking, a load on the motor; instead it is controlling one of the forward reaction steps rather than driving a reverse step. Interestingly, *f*_DNA_ can slow SMC translocation down (stopping the enzyme for large enough *f*_DNA_) but it cannot *reverse* the direction of motion of the complex down DNA in the way a load force applied directly to the enzyme might be expected to. This type of dependence of SMC translocation on *f*_DNA_ - translocation quenching but without reversal - is perhaps the clearest hallmark of this type of loop-capture model.

### C. DNA loop extrusion

Above we have shown that a loop-capture mechanism is thermodynamically capable of generating translocation of SMC complexes along a thermally fluctuating DNA molecule. It has been proposed that SMC complexes are capable of “extruding” loops of DNA, for example by the pushing apart of two motile elements [3, 47], and recent single-molecule experiments have observed such behavior [23]. We now discuss a few ways whereby the translocation model presented in Materials and Methods could be involved in active DNA loop extrusion.

#### 1. Two-translocator model

The first possibility is that two translocators of the type described in Materials and Methods could be physically linked together (Fig. 7a) so as to have a DNA loop pushed out. To have loop extrusion, the two translocators must be loaded in a specific orientation. If the DNA is topologically entrapped inside the protein “ring” of each translocator, the orientations of the two translocators will be preserved following the initial “loading”. For this model, the velocity of loop extrusion is the sum of the translocation velocities of the two translocators, or in terms of the translocation velocity of Eq. 3, *v*_loop_*_−_*_extrusion_ = 2*v*.

**FIG. 7:**
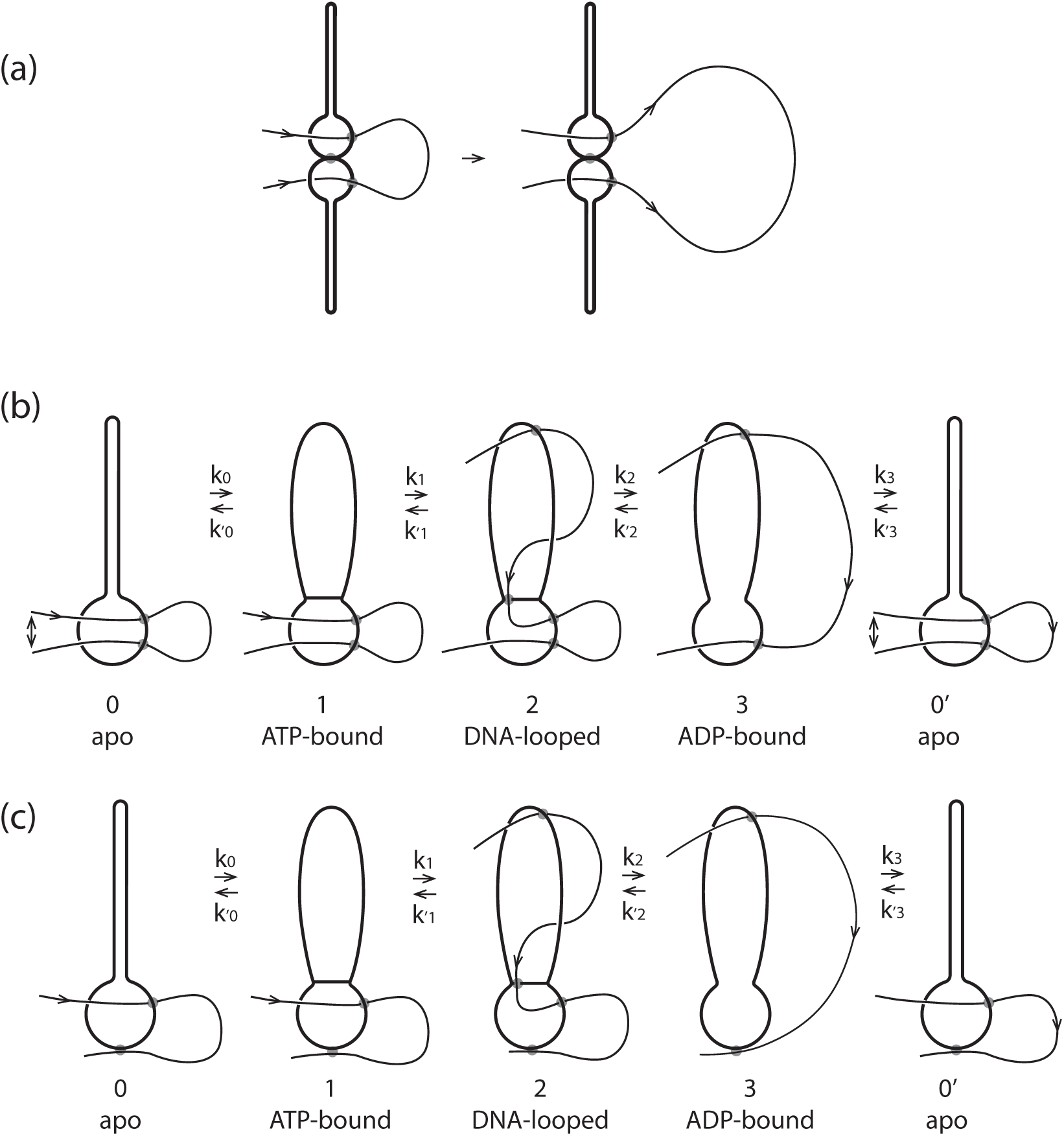
Models for loop extrusion based on the translocator model of this paper. (a) Two-translocator model. Two translocators are physically coupled together, and then loaded on DNA in orientations which generate loop growth. (b) Single-translocator model. The reaction scheme is similar to that of Fig. 4, except that two DNAs can be trapped in the lower compartment. (c) Asymmetric loop extrusion by a single translocator. The DNA remains bound to a spot outside the lower chamber (heavy black dot at bottom) resulting in one-sided loop extrusion.

#### 2. Model for loop extrusion by a single translocator

A second possibility is that a DNA loop could be extruded by a single translocator of the type described in Materials and Methods (Fig. 7b) [24]. This requires only a slight variation on the translocator model; instead of trapping just one DNA in the lower compartment (Fig. 5, state 0 and 1), we suppose that two DNAs (for instance, the base of a DNA loop) can be trapped (Fig. 7b, state 0). Starting from state 0, a small DNA loop is captured as before in the upper compartment (Fig. 7b, state 1), and then ATP hydrolysis merges the two compartments, allowing transfer of the length in the smaller (upper) DNA loop into the larger (lower) DNA loop. The reaction model for this single-translocator loop-extrusion process is just the same as for translocation (Eq. 3), *i.e.*, *v*_loop_*_−_*_extrusion_ = *v*.

The model as shown in Fig. 7b moves the upper DNA strand through the lower compartment, but the lower strand stays bound to the protein through the cycle. If iterated, this reaction will extrude a loop, but in an asymmetric manner, with one loop boundary site permanently bound. With two DNAs bound in close proximity in the lower compartment, it is likely that they will exchange binding sites periodically (likely rapidly given the high effective concentration of DNA within the lower compartment [48]), which will result in symmetric loop extrusion, still with the loop extrusion velocity equal to the DNA translocation velocity, or *v*_loop_*_−_*_extrusion_ = *v*.

#### 3. Asymmetric single-translocator model with permanently bound DNA

A variation of the single-translocator model might be one where a flanking DNA is bound to the *exterior* of the SMC protein complex through the entire reaction cycle (Fig. 7c), for example via the DNA binding sites located at the walls of the lower compartment in condensin and SMC5/6 [27–29]. If so, then translocation to the left would result in asymmetric loop extrusion, as has been observed for yeast condensin [23]. While this model provides a simple explanation for how SMC translocation can result in DNA looping, asymmetric loop extrusion is likely to perform poorly in chromosome compaction as it will tend to leave stretches of uncompacted DNA [18].

## IV. DISCUSSION

We have presented a quantitative statistical-mechanical kinetic model for SMC complexes based on structural and catalytic information for the *B. subtilis* bsSMC complex (Fig. 2). Our model follows the DNA loop-capture/protein conformational change scheme proposed in Ref. [24] (Fig. 3-5), and our theoretical analysis now shows how it can generate translocation (and loop extrusion) in reasonable accord with available experimental information. Although our model contains a number of parameters, we have chosen natural values for them, with the result that the translocation involves DNA-tension-dependent steps of from 50 nm to 150 nm in size, with a choking off of translocation for DNA tensions of *f*_DNA_ ≈ 1 pN (Fig. 6b).

### A. DNA substrate-tension *f*_DNA_ dependence of SMC translocation velocity

A key feature of our model is its dependence of translocation velocity on tension in the DNA ‘track’ that the SMC is moving along: this is unique for a molecular motor (note that myosins are not sensitive to the tension in their F-actin tracks): this reflects the essential dependence of the mechanism of our model on DNA looping, which is of course quenched by DNA tensions exceeding the pN scale [44]. Our model is an example of a novel molecular motor system, for which translocation depends on bending and consequent looping of the track itself.

This dependence of translocation on substrate tension provides an important experimental test of the model: tension in a DNA of more than a few pN should stop SMC translocation. Recently translocation of yeast condensin has been observed along DNA molecules under some tension [22], but in that experiment the applied tension was not known (however it was likely less than 1 pN). Experiments observing compaction of DNA by various types of SMC complexes have observed compaction reactions which are stalled by forces exceeding roughly 1 pN [35–38, 49], also suggestive of a loop-capture mechanism. As this paper was in preparation, quenching of loop extrusion by DNA tension ≈ 1 pN was reported for yeast condensin, providing good evidence for a DNA-loop-capture mechanism underlying DNA translocation by that enzyme complex [23].

If this relatively low (≈ 1 pN) stall force (small relative to the multi-pN stall forces of common motors like kinesins and myosins) does turn out to be a general feature of DNA-translocating SMCs, one might ask why such a “weak” motor system has been selected to play key roles in chromosome dynamics. One reason is likely that ≈ 1 pN forces are high enough to extend and transport DNA (and to bias strand transfer by type-II topoisomerases), but low enough to not significantly disturb transcription factors and DNA packaging factors non-covalently bound to the double helix [50–52]. Thus “weak” SMCs offer the feature that they can reorganize DNA without disturbing patterns of gene expression or ≈ 10 nm-scale DNA packaging machinery. Another way to look at the force scale for SMC stalling is that it is tuned so that SMCs act on DNA organization at relatively large (roughly DNA/chromatin persistence length and larger, or > 10 nm) length scales.

### B. Load force *f*_load_ and SMC translocation velocity

The substrate tension is to be distinguished from a load force applied to the SMC complex, which directly acts against translocation (Fig. 6a inset). Substrate tension is not able to push an SMC complex backwards, hence the quenching of the forward velocity (Fig. 6b). In contrast, a load force can slow the enzyme to zero velocity and even push it backwards, by forcing the enzyme to unbind from its DNA substrate (Fig. 6d).

For the parameters we have chosen, the zero-velocity point is reached at a load force of about 3.5 pN (Fig. 6d), likely dependent on details of the binding of SMC to the DNA double helix through the enzyme reaction cycle. Still, we expect the basic distinction between response to substrate tension (no reversal at large forces) and load force (reversal at large forces) shown in Figs. 6b and 6d to remain, whatever the details are of these curves.

We note that in DNA compaction assays against applied force [23, 35–38] (Fig. 8a) the applied force acts as both substrate tension *and* as a load force (the load force most likely being a fraction of the substrate tension) potentially complicating the interpretation of such experiments. Experiments of the sort of Ref. [22] studying translocation as a function of substrate tension, with a separately controlled load force (for example through attachment of a handle directly to the translocating SMC to apply a load directly) might be able to probe these two control parameters (as sketched in Fig. 6a), although applying a force to the SMC without impacting DNA tension will likely be challenging.

**FIG. 8:**
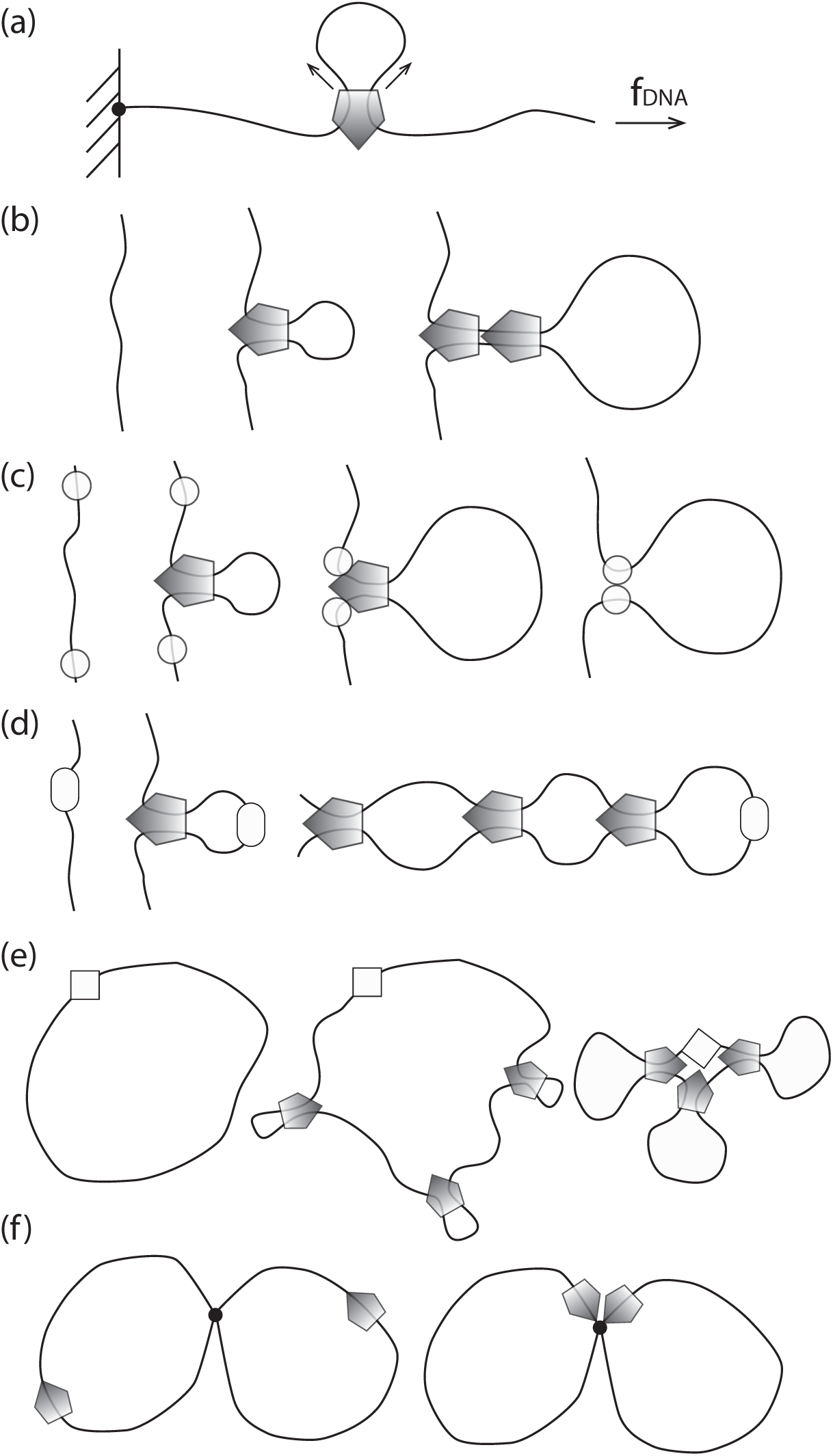
Experimental single-molecule compaction and various modes of *in vivo* SMC complex regulation. (a) Compaction assay with controlled DNA tension. The load force is related to applied tension *f* in a potentially complex way. The load force should be comparable but not larger than the tension. (b) Eukaryotic condensin: nonspecific binding and stochastic unbinding with possible ‘loop stacking’. (c) Eukaryotic cohesin: nonspecific binding (using ‘loading’ proteins) with regulated halting/unbinding. (d) Bacterial bsSMC from *B. subtilis*: site-specific binding (using the ParB protein bound to a *parS* site), with no regulation of unbinding so as to bind the two halves of a bacterial chromosome together. (e) Bacterial MukBEF from *E. coli*: site-nonspecific binding with clustering at a specific site (the origin of replication). (f) Eukaryotic SMC5/6: translocation of the complexes may lead to their co-localization at DNA junctions resulting from DNA recombination or incomplete DNA replication.

### C. Further refinements of the model

Two steps of the proposed reaction scheme may rely on a balance of SMC-DNA interactions. First, the formation of DNA loops to be captured by SMC is rate-limiting at high DNA tension in the simplest version of our DNA-segment-capture model (Fig. 6d). SMC complexes may facilitate DNA looping as well, for example by stabilizing early intermediates of the DNA looping reaction via specific SMC-DNA contacts. Such early intermediates may be captured preferentially by DNA binding at the bottom of the upper compartment, which appears to be absolutely critical for DNA translocation of bsSMC, while DNA binding at the top is not [25]. Given this, passing through the transition state may require only a reduced level of DNA bending rather than full looping, which could significantly reduce the energy barrier for the transition from states 1 to 2.

Directional translocation in our model depends on preferential capture of DNA on one side of the SMC (“upstream” in Fig. 4a), which we have argued to be a necessary result of broken symmetry of the SMC-DNA complex. While this broken symmetry is guaranteed, a strong quantitative preference for looping to one side is not, and the degree of this preference will determine the efficiency of directional translocation. Future experiments, e.g., fluorescence resonant energy transfer mapping of distances between the DNA and specific positions on the SMC, may be able to directly observe this asymmetry. Direct cryo-EM visualization may provide another way to validate this prediction of essentially any loop-capture model for SMC translocation.

Conceivably, SMC may locally ‘melt’ (strand-separate) a segment of the DNA double helix by binding single-stranded DNA and thus favour strong DNA curvatures [53] (bsSMC has long been known to bind ssDNA rather well without a clearly understood function). SMCs may also rely on other cellular proteins (such as the bacterial DNA-bending protein HU or nucleosomes) to generate strongly bent DNA *in vivo*. In our model, the quantities *φ* and *D* (see Supplementary Data) can be used to roughly take these phenomena into account, by reducing the energy barrier to loop formation (our choices of the DNA bending-energy parameter *D* and the forward/reverse rate-splitting parameter *φ* consider a completely unconstrained loop, with the effect of DNA bending free energy split equally between forward and reverse rates). For small *φ* or *D* values, the energy cost for the formation of a DNA loop is reduced, speeding up SMC translocation especially for larger DNA tensions (*>* 1 pN).

A second critical step of the model is the return of the ADP-bound state (Fig. 5, state 3) to the apo state (0) by the closure of the upper compartment and the transfer of the DNA double helix from the top of the upper compartment all the way into the lower compartment. Zipping up of the SMC arms must preferentially initiate at the top end and happen quickly so that loss of translocated DNA by free sliding through the closing upper compartment is limited. Alternatively, the DNA transfer may occur in discrete steps with DNA being fixed at each step via specific SMC-DNA contacts. We envision that this transition is critically dependent on properly folded SMC arms and blocked by SMC arms with aberrant lengths or local flexibility [41].

A less critical aspect of the model is the precise order in which, following ATP hydrolysis, ADP is released, in relation to the closing of the upper compartment (Fig. 5a, transition from state 3,2ADP to 0,2ADP). It is straightforward to reverse this order, i.e., to have ADP release precede the upper compartment closing; this will not change any of the qualitative results of the model, but will only require addition of an additional kinetic step. Similarly, the two ATP hydrolysis steps could proceed at different rates; again, this can be incorporated without any qualitative changes to the model.

### D. Application of the model to different types of SMC complexes

We have focused on bsSMC in this paper since we have more information about structural, enzymatic and *in vivo* function than any other SMC complex. We anticipate that our model will be applicable to other SMC complexes, be they eukaryote, bacterial, or archaeal. We now discuss some details of how our model might apply to specific types of SMC complexes.

#### 1. Eukaryotic condensin

Eukaryotic condensin, at present, is thought to be loaded randomly onto DNA (without specific binding sites), and then to subsequently act to compact DNA. The most simple way for this to occur is by binding, and then loop extrusion (Fig. 8b; note that the enzyme shown might be one or two SMCs of the type described in Materials and Methods. If multiple extruders bind, they can ‘stack up’ to produce more robust compacted loops [3, 18, 19].

A recent experiment has observed translocation of yeast condensin along DNA, with a velocity of roughly 100 bp/sec [22]; that experiment did not look at the translocation velocity dependence on DNA tension but it is likely that the tension in the end-tethered DNAs used was less than 1 pN, making the result consistent with our model (Fig. 6b). Single-DNA experiments studying compaction against force see a stalling of compaction for DNA tensions in the ≈ 1 to 2 pN range [35, 38], with a distribution of steps in the 50-250 nm range, also consistent with the results of our model (Fig. 6d). Additional experiments have directly observed one-sided loop extrusion, with stalling observed for DNA tensions in the pN range [23]. Future experiments might try to disentangle DNA tension and load force dependence of translocation and loop extrusion, which show somewhat different behaviors in our model.

Direct evidence for directed translocation of condensin on DNA (chromatin) *in vivo* does not yet exist, but condensin-rich structures formed at the centromeres of chromosomes in budding yeast have been argued to come about as the result of translocation of condensin on DNA[54–56].

#### 2. Eukaryotic cohesin

Eukaryotic cohesin plays at least two distinct roles in genome organization. First, it acts to hold sister chromatids together during mitosis, and is degraded by a specific protease to allow sister chromatid segregation to occur. This activity is specific for cohesin and it is unclear whether it is at all related to SMC translocation or DNA loop extrusion discussed in this paper [57].

A second, more canonical function of cohesin is to compact and organize DNA during interphase and in mitosis and meiosis, presumably via DNA loop extrusion. Cohesin is involved in the formation of topologically isolated chromosomal domains (TADs), which are thought to control gene expression at long distances by restricting or promoting enhancer/promoter interactions. In vertebrates, the two borders of a TAD are frequently defined by a pair of convergent CTCF binding sites. The precision of selection of orientation of CTCF sites at long distances observed in Hi-C experiments strongly suggests a mechanism based on processive ‘tracking’ or ‘sliding’ with CTCF acting as orientation-dependent barrier for cohesin translocation (Fig. 8c) [20, 21]. The loading of cohesin onto DNA appears to be somewhat more complicated than for condensin, and is dependent on well-characterized ‘loading factors’ which are necessary for ATP-dependent loading of cohesin onto DNA [58]. Unloading of cohesin from DNA involves another protein, Wapl [59]

Once loaded, cohesin has been observed to diffusively slide on DNA *in vitro* [60, 61] but not to undergo clear directed translocations in the way recently observed for yeast condensin. Given the long distances apparently covered by cohesin during enhancer-promoter loop formation [20, 21] there is the question of whether diffusion along DNA can transport cohesin complexes sufficiently rapidly (*i.e.* so as to reach its destination in a single cell cycle). A key open question is whether cohesin moves along DNA itself (*e.g.*, via a motor mechanism similar to that discussed in this paper), whether it undergoes directed motion by being ‘pushed’ by other DNA-motile factors (e.g., RNApol or even condensin), or whether it simply diffuses.

In conclusion cohesin is a case of an SMC complex with factors regulating its loading and unloading (Fig. 8c), with the capacity to translocate rather large distances along DNA between those events. We note that the SMC1/3 heterodimer alone has been observed to be able to trap DNA loops of 150 nm size, of a specific chirality [37], as might be expected based on our loop-trapping translocator model (Fig. 5). If cohesin proves to not itself be a DNA motor, understanding why it does not have this function, while structurally similar condensin does, will be illuminating to understand.

#### 3. Bacterial SMC complexes

This paper has relied on the large amount of structural and functional data available for the bsSMC complex from *Bacillus subtilis* [24]. Single-molecule experiments on bsSMC [49] have indicated an ATP-stimulated (but ATP-hydrolysis-independent) compaction reaction, but have not yet observed step-like translocation. The single-DNA experiments also saw appreciable DNA compaction in the absence of ATP, perhaps suggesting that bsSMC may have a simple DNA-binding-compaction function. It might be that the DNA loop-capture step of Fig. 5 (or an alternate loop-capture function) may be able to proceed in some form.

In the cell, we know that most bsSMC is initially loaded near the chromosomal origin of replication, with the help of the ParB protein which associates with the *parS* DNA sequence [16, 17, 62]. It then appears to translocate along DNA, holding the two halves (the two ‘replichores’, Fig. 8d) of the circular chromosome together [16, 17], with a translocation velocity of approximately 1 kb/sec.

The high apparent velocity of bsSMC could be achieved by the mechanism of this paper by small-loop capture at about 3 cycles per second (100 nm contains 300 bp), or perhaps with less frequent transfer of compacted DNA structures (nucleoid-associated-protein-folded DNA or small supercoiled DNA structures) through the SMC upper compartment.

The MukBEF complex supports sister chromosome segregation in *E. coli*, possibly by establishing long-range DNA contacts as other SMC complexes do. There is a long history of observations of anucleate phenotypes associated with mutations in MukBEF [63], and recent *in vivo* observations suggest that the complex operates as a dimer of SMC complexes, and then forms clusters that compact and organize the bacterial chromosome [64]. As for eukaryote condensin, no specific loading factor for MukBEF has been identified (*E. coli* lack ParB and parS), implying that DNA loop extrusion would initiate at random positions. MukBEF complexes, however, localize in sub-cellular clusters, which are usually found near the replication origins [65]. Like CTCF in the case of cohesin, replication origins might act as barriers for MukBEF translocation and thus accumulate (Fig. 8e).

Single-molecule experiments on MukBEF have observed DNA compaction against forces of up to approximately 1 pN, but without strong ATP dependence [36], rather like what has been observed for bsSMC [49]. MukBEF has been observed to be able to “cross-link” separate double helices [66]. No single-molecule experiment has yet directly observed ATP-dependent DNA compaction, ATP-dependent translocation, or ATP-dependent loop extrusion by a bacterial SMC complex.

#### 4. Eukaryotic SMC5/6

The cellular functions of SMC5/6 are related to DNA double strand break repair, the resolution of sister DNA junctions during cell division and the control of DNA topology during DNA replication. The molecular mechanisms are poorly understood and the involvement of DNA loop extrusion is unclear. The structural similarities with other SMC complexes strongly suggest shared activities, and an ATP-dependent DNA-linking function has been observed *in vitro* [67]. Conceivably, the process of DNA loop extrusion might be used to identify and locate DNA junctions for repair and resolution (Fig. 8f).

Curiously, the SMC5 and Smc6 coiled coil arms are significantly shorter than the arms of other SMC proteins [41]. Our analysis predicts that the translocation by short-arm SMC complexes might be somewhat more robust on stretched DNA. If so, then SMC5/6 complexes may have evolved to specifically deal with DNA under tension, potentially found near replication forks and during DNA repair. SMC5/6 structure may also reflect a preference for specific DNA topologies, e.g., crossings, loops or supercoiling of specific handedness.

### E. Open questions

Many basic questions about SMC complexes remain unanswered. We have proposed a model for the translocation of SMC complexes on DNA, and it is by no means clear that all SMC complexes translocate on DNA. Possibly, translocation of some SMC complexes is regulated or driven by other proteins, as may be the case for cohesin. Single-molecule approaches can provide direct observation of translocation [22] and loop-extrusion [23] behavior, and it remains an open question how general this behavior is across different species of SMCs. For loop extrusion, it is possible that multiple translocators or extruders might “stack” to provide more robust motor function [3]. How extrusion and compaction forces scale with the number of extruders stacked is also a basic and open theoretical and experimental question. At the coarse-grained level of description of the models of this paper one might ask, just what are the fundamental differences - if any - between the bacterial and eukaryote SMC complexes? For example, is there a role played by the apparent difference in symmetry between bacterial and eukaryote complexes (*e.g.*, homodimeric structure of 2x bsSMC vs. heterodimeric SMC2-SMC4 in condensin)?

## SUPPLEMENTARY DATA

Supplementary Data are available at NAR Online.

## FUNDING

This work was supported by the National Institutes of Heath [grant numbers U54-CA193419 (CR-PS-OC), U54-DK107980, and R01-GM105847 to JM], by the European Research Council [grant number CoG 724482 to SG], by the Swiss National Science Foundation [grant number 200020 163042], and by the French Agence Nationale de la Recherche [grant number ANR-14-ACHN-0016 to AB].

## Conflict of interest statement

None declared.

## Notes

#### Summary of Updates

Revision addresses referee concerns about presentation; no changes in results relative to original version. Essentially all the arithmetic has been moved to the Supplementary Data.

