## Supplementary Data for "DNA-segment-capture model for loop extrusion by structural maintenance of chromosome (SMC) protein complexes"

May 20, 2019

The supplementary data contains biophysical and mathematical details required for the definition of the model of the main text, plus the parameters used in generation of the figures. Additional figures display exploration of behavior of the model not included in the main article.

### 1 Mathematical formulation of 0↔1 transition: ATP binding and SMC head engagement

#### 1.1 Nucleotide binding kinetics

ATP association and dissociation can be described using standard binding kinetic models [1]. Starting from their apo states, the two ATP-binding sites will bind and release ATP at rates typical of ATP-binding enzymes, with an on-rate $k_{\mathrm{on,ATP}} = \gamma_{\mathrm{ATP}}[\mathrm{ATP}]$ where $\gamma$ is the reaction rate per concentration (units of $\mathrm{M}^{-1}\mathrm{s}^{-1}$) and where [ATP] is in M (mol/litre). For diffusion-limited binding reactions, $\gamma_{\mathrm{ATP}}$ has an upper limit of $\approx 10^9$ $\mathrm{M}^{-1}$ $\mathrm{s}^{-1}$ and in practice ATP-binding enzymes typically have ATP binding rates in the range of $\gamma \approx 10^6$ $\mathrm{M}^{-1}\mathrm{s}^{-1}$ [2]. The ATPs can unbind at a rate $k_{\mathrm{off,ATP}}$, which one can expect to be in the range of $10^3$ $\mathrm{s}^{-1}$ for ATPases with ATP binding site affinities ($K_{\mathrm{d,0,ATP}}$) in the mM range, such as topo II [2]. We take the dissociation constant for an ATP binding site to be $K_{\mathrm{d,0,ATP}} = k_{\mathrm{off,ATP}}/\gamma_{\mathrm{ATP}} = 0.2$ mM. Available data indicates ATP binding affinity $K_{\mathrm{d,0,ATP}} \approx$mM for bsSMC [3].

If ADP is present, it can also bind and unbind at rates $k_{\mathrm{on,ADP}} = \gamma_{\mathrm{ADP}}[\mathrm{ADP}]$ and $k_{\mathrm{off,ADP}}$; similar rates pertain for inorganic phosphate (Pi). In terms of these rates, the dissociation constant for ADP is $K_{\mathrm{d,0,ADP}} = k_{\mathrm{off,ADP}}/\gamma_{\mathrm{ADP}}$. This, and the corresponding affinity for phosphate, are likely less than those for ATP. The dashed box on the left of Fig. 5a, main text is meant to include 16 states corresponding to each Walker A site being apo, or ATP-, ADP- or ADP+Pi- bound. The total probability of all these states is $P_0$. We set $K_{\mathrm{d,0,ADP}} = K_{\mathrm{d,0,Pi}} = 20$ mM.

#### 1.2 Equilibrium probabilities of state 0 substates

In the model state 0 is actually a composite of all ATP/ADP/Pi-bound states of the SMC in conformational state 0 (Pi refers to inorganic phosphate). We treat this group of states as pre-equilibriated since the rates for ATP/ADP/Pi binding will be much faster than the adjacent SMC/DNA-conformational-mechanical transitions into SMC states 1 and 3.

The forward rate from state 0 to 1 thus includes, as a factor, the equilibrium probability of having two ATPs bound to an SMC in conformational state 0:

$$p_{0,2\mathrm{ATP}} = \left( \frac{[\mathrm{ATP}]/K_{\mathrm{d,0,ATP}}}{1 + [\mathrm{ATP}]/K_{\mathrm{d,0,ATP}} + [\mathrm{ADP}]/K_{\mathrm{d,0,ADP}} + [\mathrm{ADP}][\mathrm{Pi}]/(K_{\mathrm{d,0,ADP}} K_{\mathrm{d,0,Pi}})} \right)^2 \quad (1)$$

Similarly, the "reverse" transition from state 0 to 3 takes place through a substate which is the complex of SMC with two molecules of ADP and two molecules of inorganic phosphate, with probability:

$$p_{0,2\mathrm{ADP},2\mathrm{Pi}} = \left( \frac{[\mathrm{ADP}][\mathrm{Pi}]/(K_{\mathrm{d,0,ADP}} K_{\mathrm{d,0,Pi}})}{1 + [\mathrm{ATP}]/K_{\mathrm{d,0,ATP}} + [\mathrm{ADP}]/K_{\mathrm{d,0,ADP}} + [\mathrm{ADP}][\mathrm{Pi}]/(K_{\mathrm{d,0,ADP}} K_{\mathrm{d,0,Pi}})} \right)^2 \quad (2)$$

where $K_{\mathrm{d,0,ATP}}$ is the dissociation constant of the complex between the SMC and ATP, $K_{\mathrm{d,0,ADP}}$ is the dissociation constant of the complex between the SMC and ADP, and $K_{\mathrm{d,0,Pi}}$ is the dissociation constant of the complex between the SMC and Pi, all for SMC conformational state 0.

#### 1.3 SMC head engagement kinetics

Given pre-equilibration of nucleotide binding, the transition from the block of 0 states to the 1 state occurs at rate $k_0 = p_{0,2\mathrm{ATP}} k_{\mathrm{eng}}$, where $k_{\mathrm{eng}}$ is the rate at which engagement occurs once two ATPs are bound. The rate $k_0$ connects ATP concentration to forward cycling: $k_0$ increases linearly as [ATP] is increased from zero, and saturates at high [ATP]. We take $k_{\mathrm{eng}} = 2\ \mathrm{s}^{-1}$, *i.e.* a conformational-change time on the order of 500 msec.

We presume that the reverse head-opening rate $k_{\mathrm{dis}}$ is slower: $k_0' = k_{\mathrm{dis}} = k_{\mathrm{eng}} e^{-\beta \varepsilon_{\mathrm{eng}}}$. Here $\varepsilon_{\mathrm{eng}} = 4 k_B T$ indicates the head engagement ("sticking") free energy, since the ratio of the forward to reverse engagement rates is related to the free energy difference between engaged and disengaged states through a Boltzmann factor. This type of free energy accounting of reverse relative to forward rates will recur as we construct the reaction cycle and is essential to understanding its thermodynamics.

### 2 Mathematical formulation of 1↔2 transition: Reversible DNA loop capture

We consider loop capture by bsSMC using established mathematical models from the DNA looping and cyclization literature. The most likely size of loop $\ell$ that will form at low values of $f_{\mathrm{DNA}} < 0.1$ piconewtons (pN) will be comparable to $\ell \approx 200$ nm which is the peak of the DNA cyclization J-factor distribution [5]. At larger forces, competition between force and bending will select a loop size of $\ell = \sqrt{Dk_BTA/f_{\rm DNA}}$, where $D$ is a numerical constant equal to about 14; the energy of this loop is $\varepsilon_{\rm loop} = 2\sqrt{k_BTDAf}$ [6] which takes into account the mechanical work done against DNA tension $f_{\rm DNA}$. A suitable (free) energy and associated loop size describing the loop including the low-force limit is

$$\begin{aligned} \varepsilon_{\rm loop} &= 2\sqrt{Dk_BTA(f_0 + f_{\rm DNA})} \\ \ell &= \sqrt{Dk_BTA/(f_0 + f_{\rm DNA})} \end{aligned} \tag{3}$$

where $f_0 = 0.1$ pN provides a sensible low-force limit roughly corresponding to zero-force cyclization. As force is increased, the optimal loop size decreases, and loop energy increases. We take a fraction $\phi$ $(0 < \phi < 1)$ of the energy $\varepsilon_{\rm loop}$ as the energy barrier for the forward reaction (1→2), indicating $k_1 = k\exp[-\beta\phi\varepsilon_{\rm loop}]$. Here $k$ is a thermal fluctuation rate for an object of $d = 10$ nm length, or $k = k_BT/(\eta d^3)$; we take $\phi = 1/2$.

We need to take into account the fact that if the most probable-sized loop considered above is too small (i.e., if the force is large), it will not be able to be captured by the SMC. To include this effect we add a DNA loop-capture probability $p = 1/(1 + e^{(\ell_0-\ell)/\delta})$ with $\delta = 30$ nm and $\ell_0 = 80$ nm, to shut off the loop-capture rate when $\ell$ drops below the lower limit of loop size compatible with capture by the SMC, by more than $\delta$:

$$k_1 = kpe^{-\beta\phi\varepsilon_{\rm loop}} \tag{4}$$

Although simple, over a range of forces up to a few pN this model describes the barrier loop energy cost increase and loop size decrease as $f_{\rm DNA}$ is increased, including the inability of the SMC to capture loops which are smaller than $\ell_0$. The numerical parameter $D$ is determined by the loop bending geometry, so the precise details of the DNA conformation captured by an SMC complex can be modeled by varying the value of $D$, $\ell_0$ and $\delta$ and $\phi$.

Reversal of loop capture can occur as a result of thermal breaking of the upper-compartment DNA-SMC interaction, *i.e.*, $k_1' \propto e^{-\beta\varepsilon_{\rm bind}}$. This takes into account the binding energy of the SMC to the DNA loop, but we must also include the remainder of the loop-formation free energy which now plays the role of a tension-dependence driving the DNA loop off the complex:

$$k_1' = kpe^{-\beta\left[\varepsilon_{\rm bind} - (1-\phi)\varepsilon_{\rm loop}\right]} \tag{5}$$

where $\varepsilon_{\rm bind}$ is the energy of DNA-loop-binding; we take $\varepsilon_{\rm bind} = 15k_BT$. The bending energy ensures that the ratio of the forward and reverse rates gives the Boltzmann factor describing the relative probabilities of states 1 and 2 (with ATP bound), corresponding to a free energy difference $F_{2,{\rm ATP}} - F_1 = \varepsilon_{\rm loop} - \varepsilon_{\rm bind}$.

### 3 Mathematical formulation of 2↔3 transition: ATP hydrolysis, phosphate release, and loop release

After setting the ATP hydrolysis rate, thermodynamics dicates the synthesis rate $k_{\rm syn}$: the free energy change as one goes around the cycle of Figs. 4a and 5, main text, inferred from the net forward and reverse chemical rates must equal that released by ATP hydrolysis. The relative rates of ATP synthesis and hydrolysis must satisfy:

$$\frac{k_{\rm syn}}{k_{\rm hyd}} = \frac{K_{\rm d,2,ADP}K_{\rm d,2,Pi}}{K_{\rm d,2,ATP}}\frac{[{\rm ATP}]_{\rm eq}}{[{\rm ADP}]_{\rm eq}[{\rm Pi}]_{\rm eq}} \tag{6}$$

where the $K_{\rm d,2,X}$ refer to the binding affinities of species X to the SMC in protein state 2. These affinities are in general different from the $K_{\rm d,0,X}$ (the protein is in a different conformation) but they are likely similar in order of magnitude (the Walker A nucleotide binding site is in similar conformation).

Eq. (6) has a simple interpretation: the free energy released by ATP hydrolysis in solution (given by the log of the ratio of equilibrium concentrations) is slightly shifted by binding free energy (the log of the ratio of the affinities). Under almost any conceivable experiment conditions we will have $k_{\rm syn} \ll k_{\rm hyd}$ since the solution equilibrium of ATP is so strongly skewed towards hydrolysis, as will be shown in Sec. 5. A kinetic derivation of Eq. (6) is given in Sec. 3.2.

We take $K_{\rm d,2,X} = K_{\rm d,0,X}$ for $\rm X = ATP, ADP, Pi$ due to the lack of detailed information and also since there is relatively little effect of their precise values on the results of the model.

In the above, [Pi] and $[{\rm Pi}]_{\rm eq}$ are concentrations and equilibrium concentrations of phosphate); phosphate is involved in ATP synthesis/hydrolysis, and affects ATP/ADP free energy balance. Note that we suppose that phosphate release/binding is coincident with ADP release and binding. If phosphate release/binding is indicated by future experimental data to be at a different point in the cycle of Fig. 5a, main text, than ADP release, it is straightforward to relocate, but this will have no effect under *in-vivo*-like reactions conditions where the rate of ATP synthesis (which involves phosphate binding and therefore [Pi]) are tiny relative to that of hydrolysis.

Once hydrolysis has occurred, we presume a forward "opening" rate $k_{\rm open} = 200\ {\rm sec}^{-1}$. Given that this involves opening and relaxation of the DNA loop and release of DNA binding, we write $k_{\rm close} = k_{\rm open}e^{\beta[\varepsilon_{\rm bind}-\varepsilon_{\rm loop}-\varepsilon_{\rm open}]}$ where $\varepsilon_{\rm open} = 10k_BT$ is the additional free energy driving the SMC-DNA-ADP complex towards the open state.

#### 3.1 Elimination of ATP hydrolysis intermediates

Our model incorporates hydrolysis of two ATP molecules per SMC catalytic cycle. In steady state we can simplify the model by "integrating out" the two substates of protein state 2 with one or two ATPs hydrolysed to ADP and Pi, $[2; {\rm ATP}, {\rm ADP}\cdot{\rm Pi}]$ and $[2; 2{\rm ADP}\cdot 2{\rm Pi}]$, as follows. We will also take care to account for the inorganic phosphate Pi, which we assume to be released along with ADP. The rate equations for the various species are

$$\begin{aligned}
\frac{d[2;{\rm ATP},{\rm ADP}\cdot{\rm Pi}]}{dt} &= -(k_{\rm syn}+k_{\rm hydr})[2;{\rm ATP},{\rm ADP}\cdot{\rm Pi}] + \\
&\quad +2k_{\rm hydr}[2;2{\rm ATP}] + k_{\rm syn}[2;2{\rm ADP}\cdot{\rm Pi}] \\
\frac{d[2;2{\rm ADP}\cdot 2{\rm Pi}]}{dt} &= -(k_{\rm open}+2k_{\rm syn})[2;2{\rm ADP}\cdot 2{\rm Pi}] \\
&\quad +k_{\rm hydr}[2;{\rm ATP},{\rm ADP}\cdot{\rm Pi}] + k_{\rm close}[3,{\rm ADP}]
\end{aligned} \tag{7}$$

At steady state, the solution of these equations is

$$
\begin{aligned}
[2;\mathrm{ATP},\mathrm{ADP}\cdot\mathrm{Pi}] &= \frac{2k_{\mathrm{hydr}}(2k_{\mathrm{syn}}+k_{\mathrm{open}})}{\Delta}[2;2\mathrm{ATP}] + \frac{2k_{\mathrm{syn}}k_{\mathrm{close}}}{\Delta}[3;\mathrm{ADP}] \\
[2;2\mathrm{ADP}\cdot 2\mathrm{Pi}] &= \frac{4(k_{\mathrm{hydr}})^2}{\Delta}[2;2\mathrm{ATP}] + \frac{k_{\mathrm{close}}(k_{\mathrm{hydr}}+k_{\mathrm{syn}})}{\Delta}[3;\mathrm{ADP}]
\end{aligned}
\tag{8}
$$

where $\Delta = (2k_{\mathrm{syn}} + k_{\mathrm{open}})(k_{\mathrm{syn}} + k_{\mathrm{hydr}}) - k_{\mathrm{syn}}k_{\mathrm{hydr}}$.

By substituing these expressions in the rate equations for $[2;2\mathrm{ATP}]$ and $[3;\mathrm{ADP}]$ we find new effective transtion rates between them:

$$
\begin{aligned}
k_{2;2\mathrm{ATP}\to 3;2\mathrm{ADP}} &= k_2 = \frac{2(k_{\mathrm{hydr}})^2 k_{\mathrm{open}}}{\Delta} \\
k_{3;2\mathrm{ADP}\to 2;2\mathrm{ATP}} &= k_2' = \frac{2(k_{\mathrm{syn}})^2 k_{\mathrm{close}}}{\Delta}
\end{aligned}
\tag{9}
$$

One should keep in mind that there is still probability associated with the now "hidden" 2,ATP,ADP and 2,2ADP states

$$
\begin{aligned}
P_{2,\mathrm{ATP},\mathrm{ADP}} &= [2k_{\mathrm{hydr}}(2k_{\mathrm{syn}}+k_{\mathrm{open}})/\Delta]P_{2,2\mathrm{ATP}} \\
&+ [2k_{\mathrm{syn}}k_{\mathrm{close}}/\Delta]P_{3,2\mathrm{ADP}} \\
P_{2,2\mathrm{ADP}} &= [4(k_{\mathrm{hydr}})^2/\Delta]P_{2,2\mathrm{ATP}} \\
&+ [k_{\mathrm{close}}(k_{\mathrm{hydr}}+k_{\mathrm{syn}})/\Delta]P_{3,2\mathrm{ADP}}
\end{aligned}
\tag{10}
$$

and that $P_2 = P_{2,2\mathrm{ATP}} + P_{2,\mathrm{ATP},\mathrm{ADP}} + P_{2,2\mathrm{ADP}}$.

#### 3.2 Ratio of $k_{\mathrm{hydr}}$ to $k_{\mathrm{syn}}$

In the presence of SMCs (concentration [SMC]), a solution of ATP, ADP and inorganic phosphate Pi settles, after some time, at equilibrium concentrations dictated by the steady state of the following equations

$$
\begin{aligned}
\frac{d[\mathrm{ATP}]}{dt} &= -(k_{\mathrm{hydr},0} + k_{\mathrm{on}}^{\mathrm{SMC,ATP}}[\mathrm{SMC}])[\mathrm{ATP}] + \\
&\quad + k_{\mathrm{syn},0}[\mathrm{ADP}\cdot\mathrm{Pi}] + k_{\mathrm{off}}^{\mathrm{SMC}\cdot\mathrm{ATP}}[\mathrm{SMC}\cdot\mathrm{ATP}] \\
\frac{d[\mathrm{ADP}]}{dt} &= -(k_{\mathrm{on}}^{\mathrm{ADP,Pi}}[Pi] + k_{\mathrm{on}}^{\mathrm{SMC,ADP}}[\mathrm{SMC}])[\mathrm{ADP}] + \\
&\quad + k_{\mathrm{off}}^{\mathrm{ADP}\cdot\mathrm{Pi}}[\mathrm{ADP}\cdot\mathrm{Pi}] + k_{\mathrm{off}}^{\mathrm{SMC}\cdot\mathrm{ADP}}[\mathrm{SMC}\cdot\mathrm{ADP}] \\
\frac{d[\mathrm{Pi}]}{dt} &= -(k_{\mathrm{on}}^{\mathrm{ADP,Pi}}[\mathrm{ADP}] + k_{\mathrm{on}}^{\mathrm{SMC}\cdot\mathrm{ADP}})[\mathrm{Pi}] + \\
&\quad + k_{\mathrm{off}}^{\mathrm{ADP}\cdot\mathrm{Pi}}[\mathrm{ADP}\cdot\mathrm{Pi}] + k_{\mathrm{off}}^{\mathrm{SMC}\cdot\mathrm{ADP}\cdot\mathrm{Pi}} \\
\frac{d[\mathrm{ADP}\cdot\mathrm{Pi}]}{dt} &= -(k_{\mathrm{syn},0} + k_{\mathrm{off}}^{\mathrm{ADP}\cdot\mathrm{Pi}})[\mathrm{ADP}\cdot\mathrm{Pi}] + \\
&\quad + k_{\mathrm{hydr},0}[\mathrm{ATP}] + k_{\mathrm{on}}^{\mathrm{ADP,Pi}}[\mathrm{ADP}][\mathrm{Pi}]
\end{aligned}
$$

$$\begin{aligned}
\frac{d[\mathrm{SMC}\cdot\mathrm{ADP}]}{dt} &= -(k_{\mathrm{on}}^{\mathrm{SMC\cdot ADP,Pi}}[\mathrm{Pi}] + k_{\mathrm{off}}^{\mathrm{SMC\cdot ADP}})[\mathrm{SMC}\cdot\mathrm{ADP}] + \\
&\quad + k_{\mathrm{off}}^{\mathrm{SMC\cdot ADP\cdot Pi}}[\mathrm{SMC}\cdot\mathrm{ADP}\cdot\mathrm{Pi}] + k_{\mathrm{on}}^{\mathrm{SMC,ADP}}[\mathrm{SMC}][\mathrm{ADP}] \\
\frac{d[\mathrm{SMC}\cdot\mathrm{ADP}\cdot\mathrm{Pi}]}{dt} &= -(k_{\mathrm{off}}^{\mathrm{SMC\cdot ADP\cdot Pi}} + k_{\mathrm{syn}})[\mathrm{SMC}\cdot\mathrm{ADP}\cdot\mathrm{Pi}] + \\
&\quad + k_{\mathrm{on}}^{\mathrm{SMC\cdot ADP,Pi}}[\mathrm{Pi}][\mathrm{SMC}\cdot\mathrm{ADP}] + k_{\mathrm{hydr}}[\mathrm{SMC}\cdot\mathrm{ATP}] \\
\frac{d[\mathrm{SMC}\cdot\mathrm{ATP}]}{dt} &= -(k_{\mathrm{hydr}} + k_{\mathrm{off}}^{\mathrm{SMC\cdot ATP}})[\mathrm{SMC}\cdot\mathrm{ATP}] + \\
&\quad + k_{\mathrm{on}}^{\mathrm{SMC,ATP}}[\mathrm{SMC}][\mathrm{ATP}] + k_{\mathrm{syn}}^{\mathrm{SMC\cdot ADP\cdot Pi}}[\mathrm{SMC}\cdot\mathrm{ADP}\cdot\mathrm{Pi}] \qquad (11)
\end{aligned}$$

In these equations we capture the spontaneous hydrolysis and synthesis of ATP into and from ADP and Pi (rates $k_{\mathrm{hydr},0}$ and $k_{\mathrm{syn},0}$ respectively) alongside the same reactions catalyzed by the SMC ($k_{\mathrm{hydr}}$ and $k_{\mathrm{syn}}$). Binding/unbinding of Pi to ADP or to the SMC $\cdot$ ADP complex are explicitly taken into account.

At equilibrium, each reaction obeys detailed balance, and it is straightforward to solve for equilibrium to give an expression for the ratio of SMC-catalyzed rates: $k_{\mathrm{hydr}}/k_{\mathrm{syn}}$:

$$\frac{k_{\mathrm{hydr}}}{k_{\mathrm{syn}}} = \frac{K_{\mathrm{d,2,ATP}}}{K_{\mathrm{d,2,ADP}}K_{\mathrm{d,2,Pi}}}\frac{[\mathrm{Pi}]_{\mathrm{eq}}[\mathrm{ADP}]_{\mathrm{eq}}}{[\mathrm{ATP}]_{\mathrm{eq}}} \qquad (12)$$

Here the binding affinities are those for ATP, ADP and Pi to the SMC complex in protein state 2, each one being given by a ratio of corresponding off- to on-rates of the general form $K_{\mathrm{d}} = k_{\mathrm{off}}/k_{\mathrm{on}}$.

### 4 Energy consumption during the reaction cycle

In steady state, the net flux (total forward minus backward transitions per unit time) between each pair of adjacent states is equal, and for the four-state reduced model we have the thermodynamic relation

$$\frac{k_0k_1k_2k_3}{k_0'k_1'k_2'k_3'} = \left(\frac{[\mathrm{ATP}]/[\mathrm{ATP}]_{\mathrm{eq}}}{([\mathrm{ADP}]/[\mathrm{ADP}]_{\mathrm{eq}})([\mathrm{Pi}]/[\mathrm{Pi}]_{\mathrm{eq}})}\right)^2 \qquad (13)$$

This relation must hold since the energy dissipated during one cycle must correspond to the free energy released by hydrolysis of 2 ATPs. Eq. (13) indicates that the cycle will run "forward" (towards ATP hydrolysis) when ATP concentration is sufficiently high relative to ADP concentration, that the cycle will run backwards under conditions of sufficient excess ADP/phosphate, and that the cycle will cease when equilibrium is reached (concentrations equal to their equilibrium values). "Integrating out" the 2,ATP,ADP and 2,2ADP states does not alter the detailed balance relation (13) since $k_2/k_2' = k_{\mathrm{hyd}}^2 k_{\mathrm{open}}/(k_{\mathrm{syn}}^2 k_{\mathrm{close}})$.

The thermodynamic constraint (13) sets the as-yet-undetermined rate $k_3'$:

$$\begin{aligned}
k_3' &= k_3 p_{0,\mathrm{2ADP,2Pi}} e^{\beta[\varepsilon_{\mathrm{eng}}+\varepsilon_{\mathrm{open}}]} \\
&\quad \left(\frac{K_{\mathrm{d,2,ATP}}}{K_{\mathrm{d,0,ATP}}}\frac{K_{\mathrm{d,0,ADP}}}{K_{\mathrm{d,2,ADP}}}\frac{K_{\mathrm{d,0,Pi}}}{K_{\mathrm{d,2,Pi}}}\right)^2 \qquad (14)
\end{aligned}$$

This form of this reverse rate makes physical sense: the leading term selects only the subpopulation of state 0 with ADP and Pi bound for the transition in the direction of $k_3'$, and the Boltzmann factor provides precisely the energy cost of opening the closed "hairpin" without the ATP-closed gate. The final terms take into count the free energy difference between cofactors bound to state 0 and state 2. A explicit calculation of (13) follows.

#### 4.1 Enforcing thermodynamic consistency

One may show by explicit calculation that the physical constraint that the model dissipates only the energy associated with hydrolysis of the two ATPs per cycle, constrain the rates as described above. The basic thermodynamic relation relating the rates to energy dissipation for one cycle of the model is:

$$\frac{k_0 k_1 k_2 k_3}{k_0' k_1' k_2' k_3'} = \frac{p_{0,2\mathrm{ATP}} k_{eng} k_1 (k_{\mathrm{hydr}})^2 k_{\mathrm{open}} k_3}{k_{dis} k_1' (k_{\mathrm{syn}})^2 k_{\mathrm{close}} k_{\mathrm{rev}} p_{0,2\mathrm{ADP+2Pi}}} = \gamma \tag{15}$$

where $\gamma = 1$ in equilibrium conditions, and $\gamma = e^{2\beta\Delta G_{\mathrm{ATP}}}$ when energy is available. The hydrolysis and synthesis rates for the SMC are $k_{\mathrm{hydr}}$ and $k_{\mathrm{syn}}$. They appear squared because two ATP molecules need to be hydrolysed (or synthesized) along the cycle, and this comes naturally by using the formulas obtained in the previous section.

We note that $k_{\mathrm{rev}}$ is the transition rate from the 2ATP·2Pi substate of SMC state 0, and therefore that the transition rate from the composite state 0 is $k_3' = k_{\mathrm{rev}} p_{0,2\mathrm{ADP}\cdot 2\mathrm{Pi}}$. Similarly, the net transition rate from composite state 0 to state 1 is $k_1 = k_{\mathrm{eng}} p_{0,2\mathrm{ATP}}$.

In (15) we aim to constructively apply thermodynamic consistency to constrain the choices of rates and to derive a formula for $\gamma$.

#### 4.2 Determination of $\gamma$

We now can simply compute $\gamma$ (note that in the main text we take the approach of stating what $\gamma$ must be from energy considerations). Indeed, by substituting the expressions for $p_{0,2\mathrm{ATP}}$, $p_{0,2A\mathrm{ADP+2Pi}}$ and $k_{\mathrm{hydr}}/k_{\mathrm{syn}}$ into (15) we obtain

$$\begin{aligned} \gamma &= \frac{p_{0,2\mathrm{ATP}}}{p_{0,2\mathrm{ADP+2Pi}}} \left(\frac{k_{\mathrm{hydr}}}{k_{\mathrm{syn}}}\right)^2 \frac{k_{eng} k_1 k_{\mathrm{open}} k_3}{k_{dis} k_1' k_{\mathrm{close}} k_3'} = \\ &= \left(\frac{[\mathrm{ATP}]}{[\mathrm{ADP}][\mathrm{Pi}]} \frac{[\mathrm{ADP}]_{\mathrm{eq}}[\mathrm{Pi}]_{\mathrm{eq}}}{[\mathrm{ATP}]_{eq}}\right)^2 \frac{k_{eng} k_1 k_{\mathrm{open}} k_3}{k_{dis} k_1' k_{\mathrm{close}} k_{\mathrm{rev}}} \end{aligned} \tag{16}$$

By evaluating this expression at equilibrium ($[\mathrm{ATP}] = [\mathrm{ATP}]_{eq}$, $[\mathrm{ADP}] = [\mathrm{ADP}]_{eq}$, $[\mathrm{Pi}] = [\mathrm{Pi}]_{\mathrm{eq}}$ and $\gamma = 1$) we obtain

$$\frac{k_{eng} k_1 k_{\mathrm{open}} k_3}{k_{dis} k_1' k_{\mathrm{close}} k_{\mathrm{rev}}} = 1 \tag{17}$$

and thus we have

$$\gamma = \left(\frac{[\mathrm{ATP}]}{[\mathrm{ATP}]_{eq}} \frac{[\mathrm{ADP}]_{eq}}{[\mathrm{ADP}]} \frac{[\mathrm{Pi}]_{eq}}{[\mathrm{Pi}]}\right)^2 \tag{18}$$

In this expression we still need the phosphate concentrations, [Pi] and $[\mathrm{Pi}]_{\mathrm{eq}}$. In biochemistry experiments, one typically strives for large excess of ATP over Pi, and it is often difficult to say what the Pi concentration is for in vitro experiments. Fortunately, in a driven system that primarily runs "forward", the synthesis reaction will often be so slow that one can choose a Pi concentration with impunity. Alternately, in vivo, $[\mathrm{Pi}] \simeq 20-40$ mM [7], implying that, Pi in excess over the nucleotides (which in vivo are in the mM range) and thus will be unaffected by ATP hydrolysis/synthesis, allowing the approximation $[\mathrm{Pi}] = [Pi]_{eq}$, simplifying in the expression for $\gamma$.

##### 4.3 Reaction cycle flux

The complete formula for the flux (steady state reaction cycles per unit time) is too complex to publish: the authors will of course provide the formula on request. However, it is possible to show analytically that it has the form

$$\mathrm{flux} = (\gamma - 1)\, k_1 k_3 k_{eng} k_{\mathrm{hydr}}^2 k_{\mathrm{open}} F\left([\mathrm{ANP}], \frac{[\mathrm{ATP}]}{[\mathrm{ADP}]}, \frac{[\mathrm{ATP}]_{\mathrm{eq}}}{[\mathrm{ADP}]_{\mathrm{eq}}}\right) \tag{19}$$

The $\gamma - 1$ factor in this expression shows that the flux is proportional to the exponential of the energy available from ATP hydrolysis. This property verifies that the model is thermodynamically consistent.

#### 5 Free energy release from ATP hydrolysis and equilibrium nucleotide concentrations

One can consider as input to the model the concentrations of phosphate and nucleotides, i.e., [ATP], [ADP] and [Pi] as prepared in a given experimental situation or as occur *in vivo*. Our model also requires the equilibrium concentrations of these reactants given those inputs, which is determined by the dissociation constant $K$ for ADP/ATP equilibrium, via $[\mathrm{ADP}]_{\mathrm{eq}}[\mathrm{Pi}]_{\mathrm{eq}} = K[\mathrm{ATP}]_{\mathrm{eq}}$. Experimental data indicate $K = e^{16}$ M [8]; given that in vitro or in vivo [ATP] is far above its equilibrium value allows use of the first term of an expansion in $[\mathrm{ATP}]/K$, of the exact result for $[\mathrm{ATP}]_{\mathrm{eq}}$ (derived in Sec. 5.1):

$$[\mathrm{ATP}]_{\mathrm{eq}} = \frac{([\mathrm{ADP}] + [\mathrm{ATP}])\,([\mathrm{Pi}] + [\mathrm{ATP}])}{K} + \mathcal{O}\left(\frac{[\mathrm{ANP}]^3}{K^2}\right) \tag{20}$$

with $[\mathrm{ANP}] = [\mathrm{ADP}] + [\mathrm{ATP}]$, $[\mathrm{ADP}]_{\mathrm{eq}} = [\mathrm{ADP}] + [\mathrm{ATP}] - [\mathrm{ATP}]_{\mathrm{eq}}$ and $[\mathrm{Pi}]_{\mathrm{eq}} = [\mathrm{Pi}] + [\mathrm{ATP}] - [\mathrm{ATP}]_{\mathrm{eq}}$.

##### 5.1 Equilibrium ATP, ADP and Pi concentrations

Since one ADP and one Pi can combine to form one ATP, the *total* ADP and Pi including that "bound" inside ATP are $[\mathrm{ADP}]_{\mathrm{T}} = [\mathrm{ADP}] + [\mathrm{ATP}]$ and $[\mathrm{Pi}]_{\mathrm{T}} = [\mathrm{Pi}] + [\mathrm{ATP}]$. The equilibrium concentrations are therefore related by $([\mathrm{ADP}]_{\mathrm{T}} - [\mathrm{ATP}]_{\mathrm{eq}})([\mathrm{Pi}]_{\mathrm{T}} - [\mathrm{ATP}]_{\mathrm{eq}}) = K[\mathrm{ATP}]_{\mathrm{eq}}$, where $K$ is the equilibrium constant for the ATP synthesis reaction.

We may solve for $[\text{ATP}]_{\text{eq}}$:

$$
\begin{aligned}
[\text{ATP}]_{\text{eq}} &= \frac{1}{2}\Big[K + [\text{ADP}] + [\text{Pi}] + 2[\text{ATP}] \\
&- \sqrt{(K + [\text{ADP}] + [\text{Pi}] + 2[\text{ATP}])^2 - 4([\text{ADP}] + [\text{ATP}])([\text{Pi}] + [\text{ATP}])}\Big] \qquad (21)
\end{aligned}
$$

with $[\text{ADP}]_{\text{eq}} = [\text{ADP}] + [\text{ATP}] - [\text{ATP}]_{\text{eq}}$ and $[\text{Pi}]_{\text{eq}} = [\text{Pi}] + [\text{ATP}] - [\text{ATP}]_{\text{eq}}$.

Given that the decomposition is ATP is highly exothermic under most circumstances, $K$ will be large compared to any of the reactant concentrations, indicating that we can expand (21) in powers of $1/K$:

$$
[\text{ATP}]_{\text{eq}} = \frac{([\text{ADP}] + [\text{ATP}])\,([\text{Pi}] + [\text{ATP}])}{K} + \mathcal{O}\left(\frac{[\text{ANP}]^3}{K^2}\right) \qquad (22)
$$

Experimental data indicate $[\text{ATP}]_{\text{eq}}/[\text{ADP}]_{\text{eq}} \approx 10^{-9}$ under *in vivo*-like conditions with $[\text{Pi}] \approx$ 10 mM [8], indicating $K = e^{16}$ M, the value taken in the main paper.

### 6 Michaelis-Menten-like cycling rate for the approximate model

Starting from the reduced 4-state model, we neglect all reverse rates except $k_1'$ and setting $P_2 = P_{2,\text{ATP}}$, the steady-state equations simplify to:

$$
\begin{aligned}
k_3 P_3 &= k_0 P_0 \\
k_0 P_0 + k_1' P_2 &= k_1 P_1 \\
k_1 P_1 &= (k_1' + k_2) P_2 \\
k_2 P_2 &= k_3 P_3 \qquad (23)
\end{aligned}
$$

As for Eq. 13 of the main paper, these equations are overdetermined, and any one can be dropped. The resulting inhomogenous linear system of four equations for the four $P_i$ can easily be solved, with the result

$$
\begin{aligned}
P_0 &= \frac{k_1 k_2 k_3}{k_1 k_2 k_3 + (k_1 k_2 + k_1 k_3 + k_1' k_3 + k_2 k_3) k_0} \\
P_1 &= \frac{[k_1' + k_2] k_0}{k_1 k_2} P_0 \\
P_2 &= \frac{k_0}{k_2} P_0 \\
P_3 &= \frac{k_0}{k_3} P_0 \qquad (24)
\end{aligned}
$$

The steady-state cycling rate can be computed from any one of the transition steps in the model. Because of the irreversibility of three of the steps (head engagement, ATP hydrolysis,

ADP release), the reaction can only cycle forward; the cycling rate $k_{\text{cycle}}$ can be most easily be computed from one of the irreversible steps, *e.g.*, $k_{\text{cycle}} = k_0 P_0$, or

$$\begin{aligned} k_{\text{cycle}} &= \frac{k_1 k_2 k_3 k_0}{k_1 k_2 k_3 + (k_1 k_2 + k_1 k_3 + k_1' k_3 + k_2 k_3) k_0} \\ &= \frac{k_0}{1 + (B + C/k_1) k_0} \end{aligned} \tag{25}$$

where $B = 1/k_2 + 1/k_3$ and $C = 1 + k_1'/k_2$ are used to write the simplified final expression.

### 7 Additional results for the model

We include a series of figures showing the steady-state translocation/extrusion rate (ignoring any slippage effects from load force) and state probabilites calculated for the reduced 4-state model with all four reverse rates (i.e., not the approximate model of the previous section), which may be useful for reproducing our results.

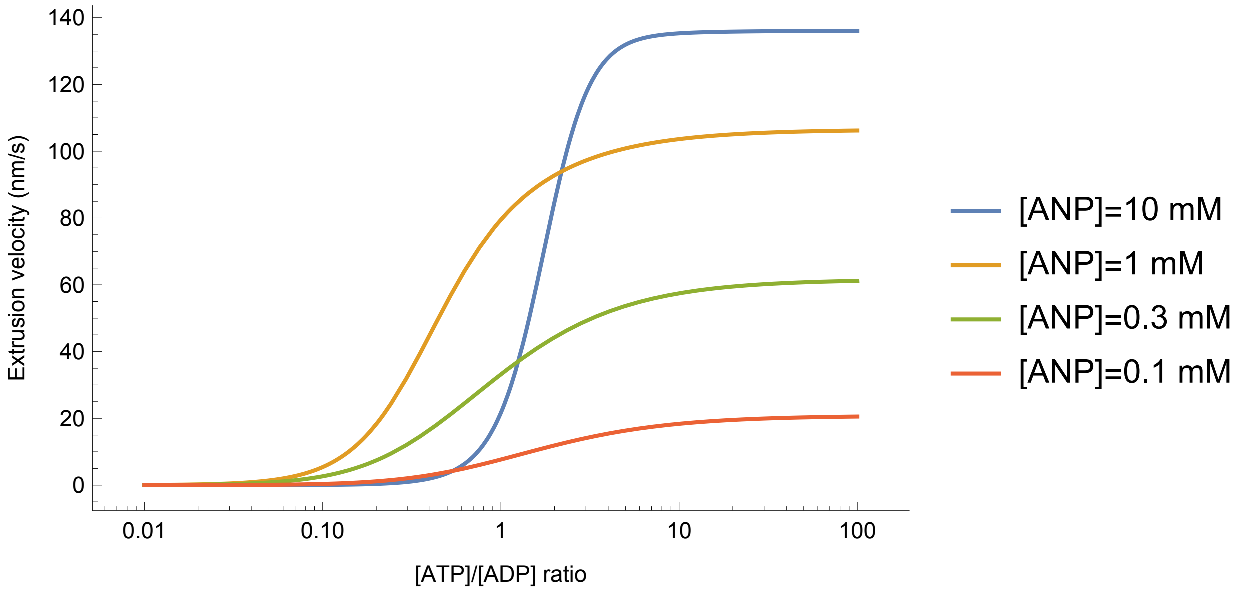


Supplementary Figure 1: Extrusion/translocation velocity as a function of the [ATP]/[ADP] ratio for various total nucleotide concentrations.

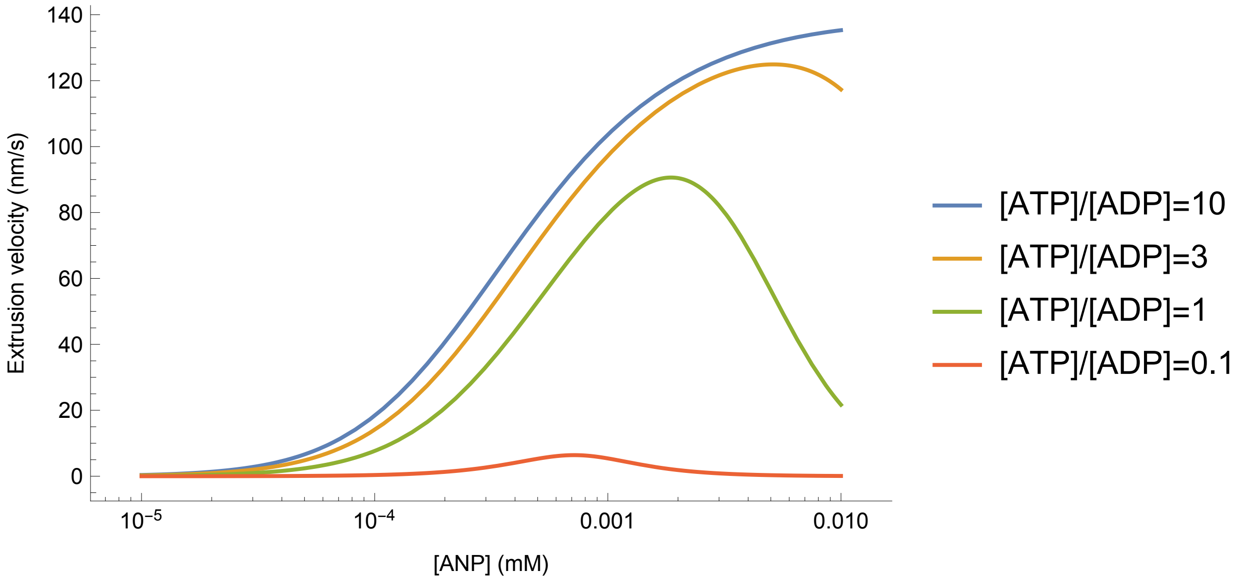


Supplementary Figure 2: Extrusion/translocation velocity as a function of the total nucleotide concentration, [ANP], for various [ATP]/[ADP] ratios.

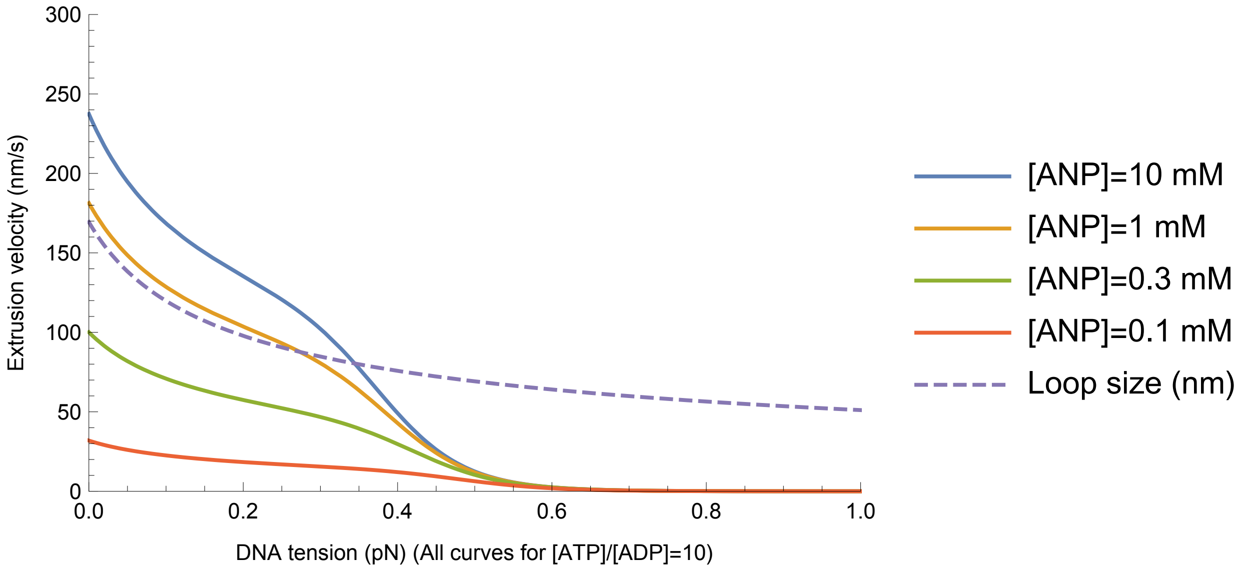


Supplementary Figure 3: Extrusion/translocation velocity as a function of the DNA tension, for a series of fixed total nucleotide concentration, [ANP]=[ATP]+[ADP].

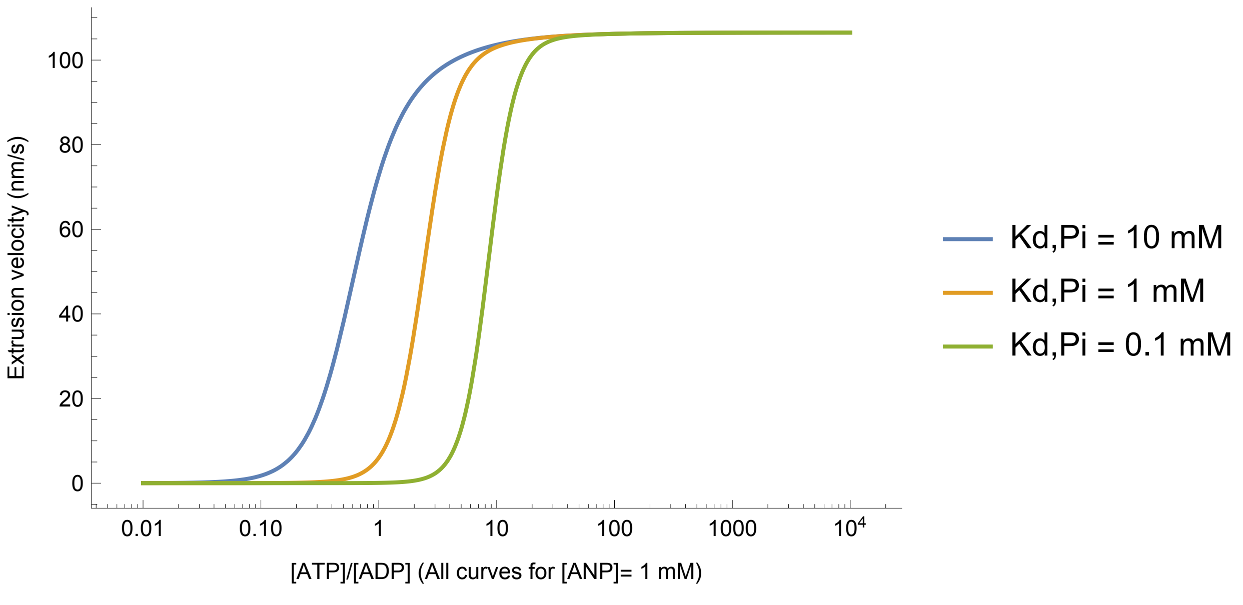


Supplementary Figure 4: Extrusion/translocation velocity as a function of the [ATP]/[ADP] ratio for various phosphate dissociation constants (and [ANP] = 1 mM). It can be observed that, the stickier is phosphate, the larger the [ATP]/[ADP] ratio must be to have extrusion. This can be understood thermodynamically, because more energy (proportional to $\ln([ATP]/[ADP])$ is consumed to detach the phosphate, and less is available to extrude loops.

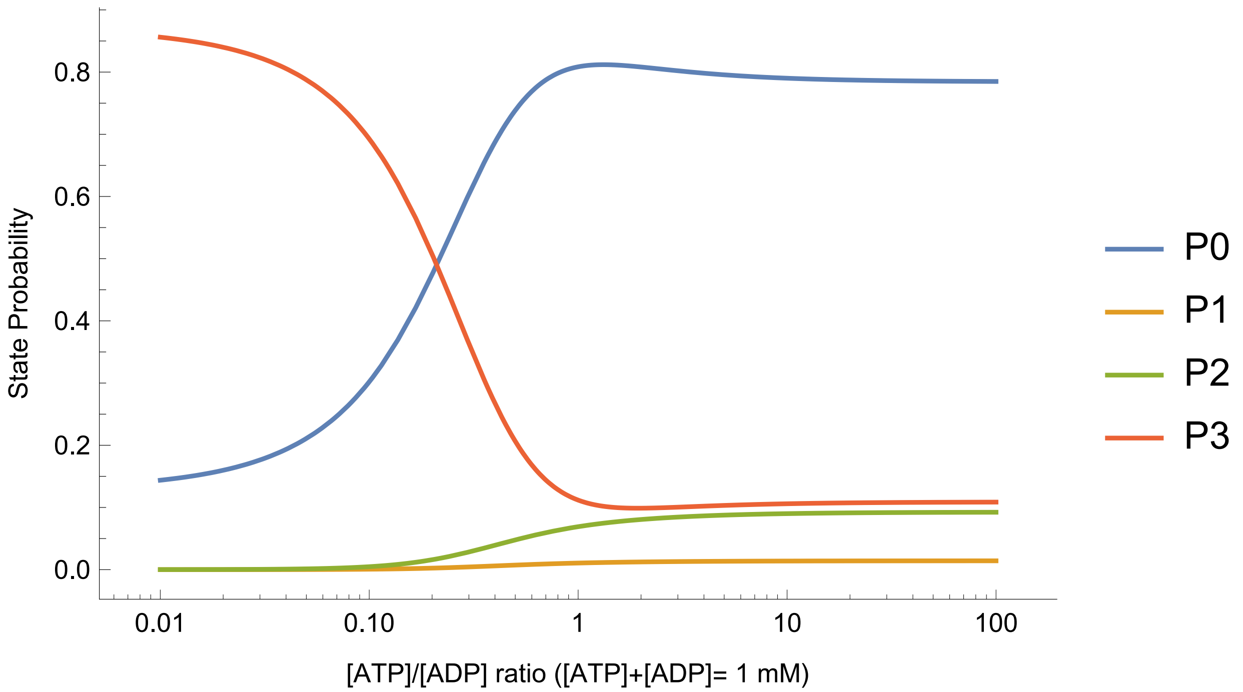


Supplementary Figure 5: Probabilities of the states of the reduced model as a function of the [ATP]/[ADP] ratio ([ANP = 1 mM]). $P_2$ refers to the sum of probabilities of the three nucleotide occupation states in SMC state 2.

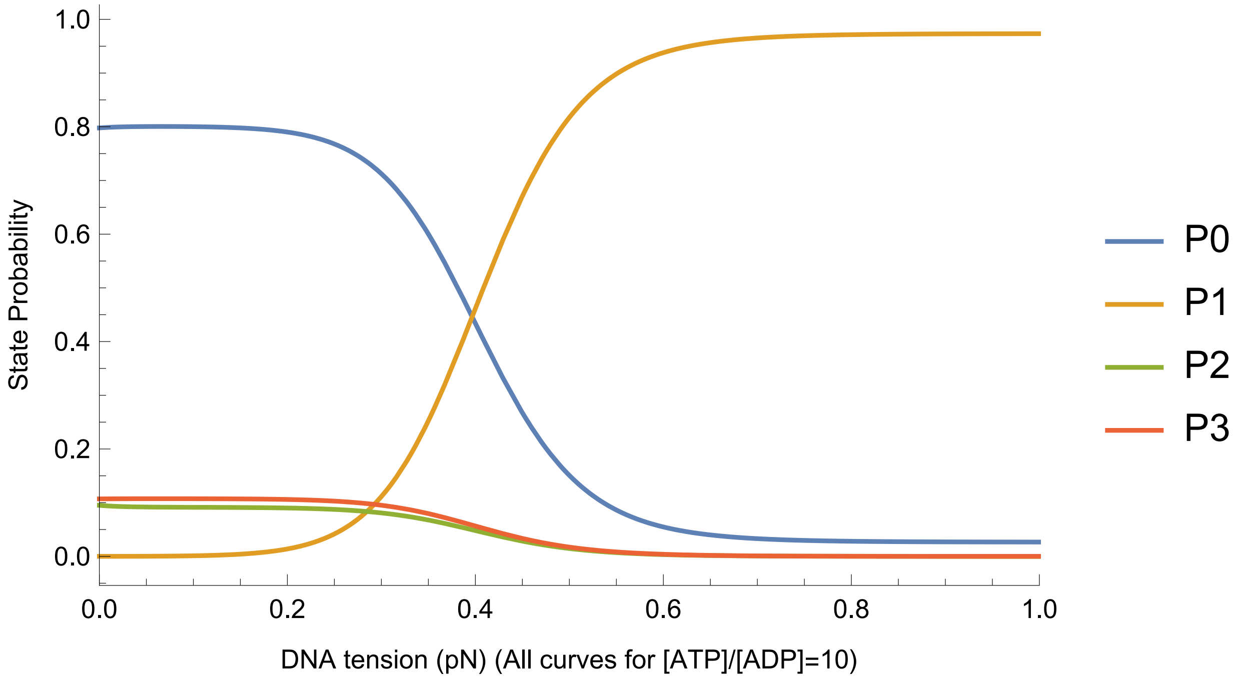


Supplementary Figure 6: Probabilities of the states of the reduced model as a function of the DNA tension for [ATP]/[ADP]=10 and[ANP] = 10 mM]. $P_2$ refers to the sum of probabilities of the three nucleotide occupation states in SMC state 2.
